# GLABRA2 regulates gene expression via its own EAR-motif mediated recruitment of the TPL/TPR corepressors

**DOI:** 10.64898/2026.08.26.747311

**Authors:** Bilal Ahmad, Aytug Ulutas, Ayianna K. Bailey, Lillian R. Marberg, Kathrin Schrick

**Affiliations:** Division of Biology, Kansas State University, Manhattan, KS 66506, USA; Interdepartmental Genetics Program, Kansas State University, Manhattan, KS 66506, USA

**Keywords:** HD-Zip IV transcription factors, Arabidopsis, repression, EAR motif, GL2

## Abstract

The Arabidopsis HD-Zip IV transcription factor GLABRA2 (GL2) displays dual regulatory capabilities, as an activator and repressor of genes that mediate cell-type differentiation of the epidermis. GL2 binds L1 box elements in the promoters of its target genes; however, the mechanisms by which it controls gene expression remain elusive. GL2 contains two putative ethylene-responsive element-binding factor-associated amphiphilic repression (EAR) motifs proximal to its N– and C-termini. The N-terminal EAR motif is highly conserved among GL2 orthologs that form a distinct clade of HD-Zip IV transcription factors in monocots and dicots. We demonstrate that deletion or Ala substitution of this N-terminal EAR motif results in a partial loss-of-function phenotypes in trichomes, non-hair root cells, and seed coat mucilage. In contrast, mutations affecting the C-terminal EAR motif display improper nuclear localization, likely due to protein misfolding. Yeast two-hybrid and in planta co-immunoprecipitation assays show that GL2 selectively interacts with the TOPLESS (TPL) and TPL-RELATED (TPR) corepressors via its N-terminal EAR motif. Fusion of the SUPERMAN REPRESSIVE DOMAIN X (SRDX) with the *gl2* N-terminal EAR motif mutant (*gl2^EAR-N^*) rescues the epidermal defects of *gl2* mutants. Transcriptome analysis of mutant and wild-type seedling roots further confirms the role of the GL2 N-terminal EAR motif in tuning gene expression. Our findings support a model whereby GL2 recruits TPL/TPR corepressors via its EAR motif to sequester histone-modifying proteins, resulting in chromatin remodeling required for epidermal development.

## Introduction

GLABRA2 (GL2) represents a distinct clade of the class IV homeodomain leucine-zipper (HD-Zip IV) family, and was initially discovered three decades ago for its role in trichome differentiation (Rerie et al., 1994; Chew et al., 2013). Subsequent studies revealed its role in the regulation of ectopic root hair formation (Di Cristina et al., 1996; Masucci et al., 1996), seed coat mucilage synthesis (Western et al., 2001), seed oil accumulation (Shen et al., 2006; Shi et al., 2012), and anthocyanin biosynthesis (Wang et al., 2015). GL2 is additionally implicated in stress responses and nutrient starvation (Cheng et al., 2024; Navarro et al., 2024). Under nutrient-deficient conditions, GL2 expression decreases, enhancing ethylene production and root elongation (Cheng et al., 2024). Furthermore, GL2, in concert with ANTHOCYANINLESS2 (ANL2), controls arsenic signaling and accumulation (Navarro et al., 2024). Under low arsenic conditions, GL2 binds to the promoter of *ASK18*, essential for arsenic detoxification and stress tolerance, resulting in its transcriptional repression (Navarro et al., 2024).

GL2 orthologs and other HD-Zip IV family members have been characterized in various crops, establishing their significance for plant growth and development (Guan et al., 2008; Vernoud et al., 2009; Chew et al., 2013; Wu et al., 2023). GL2-type *RICE OUTERMOST CELL-SPECIFIC* (*ROC*) genes in rice exhibit predominant expression in epidermal tissues (Ito et al., 2002; Ito et al., 2003; Chew et al., 2013). Among these, *ROC5* and *ROC8* regulate bulliform cell formation and leaf rolling, whereas *ROC4* mediates drought tolerance by controlling wax synthesis (Zou et al., 2011; Wang et al., 2018; Xu et al., 2021). *ROC4* and *ROC5* contribute to drought stress tolerance and resistance against brown planthopper infestation by modulating cuticle wax production and bulliform cell formation (Tao et al., 2024). identified Similarly, a maize *GL2* ortholog *OUTER CELL LAYER* 4 (*ZmOCL4*) plays an essential role in regulating trichomes and anther development (Vernoud et al., 2009). Cotton *HOMEOBOX GENE 1* (*GaHOX1*), a functional ortholog of *GL2*, rescues epidermal defects in *gl2* null mutants when expressed under the *GL2* promoter in Arabidopsis (Guan et al., 2008). *GaHOX1 and GaHOX3* drive the development of cotton fiber (Guan et al., 2008; Shan et al., 2014). *Woolly*, a tomato ortholog of *GL2*, dose-dependently regulates the development and morphology of trichomes (Wu et al., 2023).

Like other eukaryotic transcription factors, GL2 serves a dual role as a transcriptional repressor and activator. We previously demonstrated that GL2 functions as a transcriptional activator to promote the expression of the R2R3-MYB transcription factor *MYB23* through a positive feedback loop (Khosla et al., 2014). Wang et al. reported that GL2 binds the *VILLIN 1 (VLN1)* promoter to positively regulate its expression in response to osmotic stress in root hairs (Wang et al., 2020b). In contrast, Ohashi et al. showed that GL2 negatively regulates the expression of *PHOSPHOLIPASE Dz1* (*PLDz1*), in non-hair cell files (Ohashi et al., 2003). In another example of is repressive activities, GL2 negatively regulates basic helix-loop-helix (bHLH) transcription factor genes, including *ROOT HAIR DEFECTIVE6 (RHD6), RHD6-LIKE1 (RSL1), RSL2, Lj-RHL1-LIKE1 (LRL1)*, and *LRL2* (Lin et al., 2015). The functional dichotomy of GL2 may be due to its interaction with specific corepressor proteins that operate spatially and/or temporally.

The plant corepressors, TPL and TPR1-4, are named for the distinctive “topless” shoot developmental defects observed in the *tpl-1* mutant (Long et al., 2002; Long et al., 2006). TPL and TPR proteins are recruited by transcription factors containing a repression domain (RD), which facilitates this interaction. This recruitment can occur through direct binding with an RD-containing transcription factor or adaptor protein that links the corepressor to DNA binding proteins (Plant et al., 2021). The ethylene-responsive element-binding factor-associated amphiphilic repression (EAR) motif, a specific type of repression domain (RD), was initially identified in class II ethylene response factor (ERF) proteins of *Nicotiana tabacum* (Ohta et al., 2001). This motif is present in approximately 12% of transcriptional repressor proteins in Arabidopsis (Plant et al., 2021). It is characterized by a consensus sequence of (L/F)DLN(L/F)xP or LxLxL and is responsible for recruiting the corepressors (Kagale et al., 2010). TPL/TPR proteins interact with histone deacetylases such as HDA6 and HDA19, leading to histone hypoacetylation and subsequent transcriptional repression of target loci (Zhu et al., 2010; Wang et al., 2013; Liu et al., 2014; Oh et al., 2014).

A subset of EAR motif-containing proteins also interact with SWI-INDEPENDENT 3-ASSOCIATED POLYPEPTIDE18 (SAP18), another corepressor protein (Song and Galbraith, 2006; Hill et al., 2008; Kagale and Rozwadowski, 2011). SAP18 serves as a component of two distinct complexes: SIN3-HISTONE DEACETYLASE COMPLEXES (SIN3/HDACs) and APOPTOSIS AND SPLICING-ASSOCIATED PROTEIN (ASAP) complexes (Zhang et al., 1997; Deka and Singh, 2017). SAP18 also co-purifies with HDA19 and various polycomb repressive complex 2 (PRC2) core components (Qüesta et al., 2016). The PRC2 components CURLY LEAF (CLF) and SWINGER (SWN) function as methyltransferases that catalyze the trimethylation of histone H3 at Lys 27 (H3K27me3), an epigenetic modification associated with transcriptional repression, acting either independently or in synergy with histone deacetylation (Shu et al., 2019; Baile et al., 2021; Bieluszewski et al., 2021; Cai et al., 2021).

Here we address mechanism of GL2-mediated gene repression. A previous study suggested that GL2 interacts with two EAR-motif-containing adaptor proteins, GL2-INTERACTING REPRESSOR 1 (GIR1) and GIR2 (Wu and Citovsky, 2017a). This interaction was reported to require TPL recruitment, leading to repression of GL2 targets in roots (Wu and Citovsky, 2017b). Our findings contradict this earlier report and reveal that GL2 itself contains an N-terminal EAR motif, which is conserved among GL2 orthologs and is essential for transcription factor activity. Leveraging deletion and substitution mutational analyses, we demonstrate that the GL2 N-terminal EAR motif is crucial for normal epidermal development in trichomes, roots and seeds. Protein-protein interaction analysis, utilizing complementary approaches, indicates that GL2 physically interacts with TPL and TPR corepressors. N-terminal fusion of the synthetic EAR motif SUPERMAN REPRESSIVE DOMAIN X (SRDX) restored function, providing evidence that the EAR motif is sufficient for GL2 activity as a transcriptional repressor. We propose a model whereby GL2 requires an EAR motif to interact with TPL/TPR corepressor proteins which subsequently recruit chromatin modifiers, leading to the repression of target genes driving epidermal cell-type specification.

## Results

### GL2 acts independently of GIR1, an EAR-motif containing adaptor protein that physically interacts with a subset of HD-Zip IV transcription factors

The Arabidopsis *GIR1/At5g06270* and *GIR2/At3g11600* genes, which share a high degree of sequence similarity, were previously described to encode adaptor proteins that physically interact with both GL2 (Wu and Citovsky, 2017a) and TPL (Wu and Citovsky, 2017b). Due to their presumed role in conjunction with *GL2*, these genes were named *GL2-INTERACTING REPRESSOR1* and *2* (*GIR1* and *GIR2*). To test for genetic interactions between *GL2* and *GIR1*, we constructed the double mutants. Surprisingly, the *gl2-5;gir1* mutants display enhanced trichome formation on leaves, indicating that the *gir1* mutation can lead to extragenic suppression of *gl2-5,* a null mutant allele (**Fig. S1**). Importantly, this result implies that the *GIR1* gene does not require GL2 for its activity in mediated trichome cell differentiation. Crosses to *hdg11* and analysis of *gl2;gir1;hdg11* triple mutants revealed that the enhanced trichome formation in *gl2;gir1* double mutants is likely due to upregulation of *HDG11*, another HD-Zip IV transcription factor family member. Intriguingly, the *hdg11-1;gl2-5;gir1* triple mutants exhibit similar trichome phenotypes as *hdg11-1;gl2-5* double mutants (**Fig. S1**), consistent with the idea that *GIR1* acts through *HDG11,* but not *GL2*.

To check the reported physical interaction between GIR1 and GL2 (Wu and Citovsky, 2017a), we performed yeast two-hybrid (Y2H) assays with the full-length proteins. Contrary the previous report, our results demonstrate that GIR1 and GIR2 fail to interact with GL2 when tested in bait or prey combinations (**Fig. S2A**). Instead, the Y2H data show that GIR1 strongly interacts with a subset of HD-Zip IV transcription factors, including HDG11, ATML1, PDF2, HDG1, HDG2, and HDG7 (**Fig. S2B**). Overall, our findings suggest diversification of the HD-Zip IV transcription factor family, and corepressor binding activity that is independent of the GIR1 adaptor protein. We hypothesized that GL2 and its orthologs interact directly with corepressor proteins.

### N-terminal EAR motif is conserved among GL2 orthologs in monocots and dicots

Genome-wide prediction of EAR motif-containing proteins identified the consensus sequence (_22_LSLSL_26_) within the N-terminus of GL2 (Kagale et al., 2010). Our analysis uncovered another putative EAR motif at the C-terminus (_703_LTLAL_707_) (**Fig. 1A**). Both motifs are predicted in PlantEAR (http://structuralbiology.cau.edu.cn/plantEAR/), a functional analysis platform that incorporates the Hidden Markov Model and orthologous gene searches across 71 plant species (Yang et al., 2018). In Arabidopsis, a subset of HD-Zip IV members (7 out of 16) contains a LxLxL type motif, which varies in position (**Table S1**). Notably, GL2 is the only Arabidopsis HD-Zip IV member with two putative EAR motifs, while HD-Zip III members are not predicted to contain an EAR motif. Multiple sequence alignments of GL2 orthologs from various monocots and dicots revealed that the N-terminal EAR motif is highly conserved (**Fig. S3**). In contrast, the putative C-terminal EAR motif exhibited less conservation that was limited to closely related Brassicaceae species including *Camelina sativa*, *Capsella rubella*, and *Brassica napus*.

**Figure 1.**
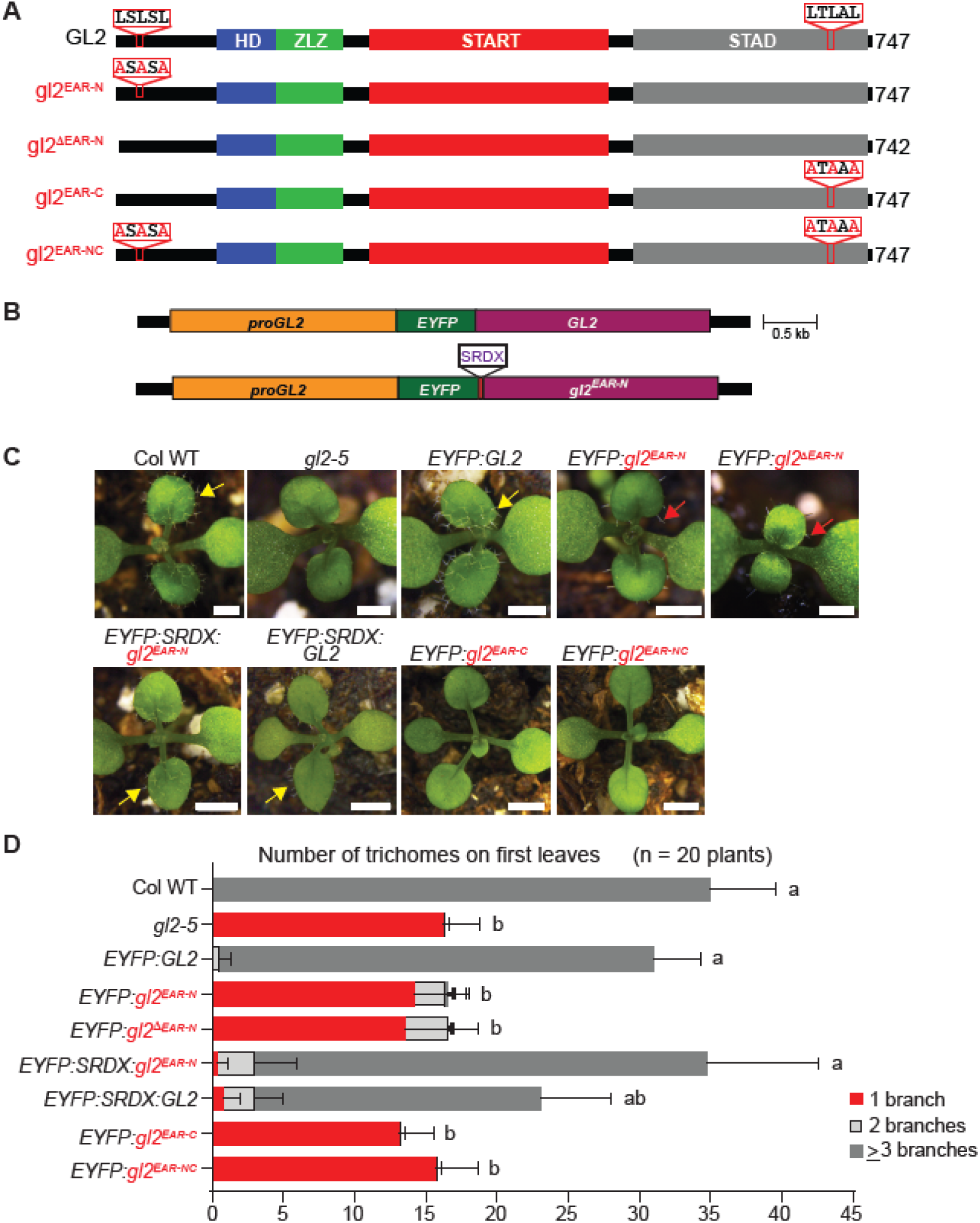
The *gl2* EAR motif mutants exhibit trichome defects. (A) Schematics indicate domain configurations of wild-type GL2 and EAR motif mutants. HD, Homeodomain; ZLZ, Zipper Loop Zipper; START, Steroidogenic Acute Regulatory (StAR) related lipid Transfer; STAD, START Adjacent Domain. EAR motifs and positions of Leu to Ala substitution mutants are shown. Amino acid lengths are given on the right. (B) Schematic of transgenic constructs for EYFP-tagged *GL2* and *SRDX* fused to the *gl2* N-terminal EAR-motif mutant (*gl2^EAR-N^*). The native *GL2* promoter (*proGL2*) drives gene expression of the transgenes. (C) Trichome phenotypes on first leaves. Columbia wild type (Col WT) and *gl2-*5 plants transgenic for *EYFP:GL2*, *EYFP:SRDX:GL2*, and *EYFP:SRDX:gl2^EAR-N^* exhibit normal trichome formation (yellow arrow), while the *gl2-5* null mutant, and the *gl2^EAR-C^* and *gl2^EAR-NC^* EAR-motif mutants display trichome defects. The *gl2^EAR-N^ and gl2^ΔEAR-N^* EAR-motif mutants exhibit a partial trichome phenotype (red arrow). Bar = 1 mm. (D) Quantification of trichome numbers and branching on first leaves for the genotypes shown in **(C)**. Error bars depict the SD, and letters denote significant between genotypes as determined by one-way ANOVA and Tukey’s test (p < 10^-5^).

### N-terminal EAR motif of GL2 is required for epidermal differentiation in the leaf, root, and seed

To examine the function of the putative EAR motifs, we generated an N-terminal Ala substitution mutant (*gl2^EAR-N^*), an N-terminal EAR motif deletion mutant (*gl2^ΔEAR-N^*), a C-terminal Ala substitution mutant (*gl2^EAR-C^*), as well as a double N-and C-terminal Ala substitution mutant (*gl2^EAR-NC^*) (**Fig. 1A**). Wild-type and mutant coding sequences were fused with *EYFP* and expressed under the native *GL2* promoter (**Fig. 1B**). The resulting constructs were transformed into the *gl2-5* null mutant background to obtain stable homozygous lines. The untransformed *gl2-5* null mutant and wild type served as negative and positive controls, respectively. The *proGL2:EYFP:GL2* lines displayed rescue of the *gl2-5* phenotype, displaying normal trichome formation like that of the wild type, whereas the N-terminal EAR motif mutants (*gl2^ΔEAR-N^*, *gl2^EAR-N^*) displayed a partial trichome phenotype (**Fig. 1C**). In contrast, the *gl2^EAR-C^* and *gl2^EAR-NC^* mutants exhibited glabrous leaves like that of *gl2-5* (**Fig. 1C**). Quantification of trichomes and branching patterns confirmed that the N-terminal EAR motif mutants exhibit significant increases in 2– or 3-branched trichomes, a partial function phenotype in comparison to the null mutant, whereas the *gl2^EAR-C^* and *gl2^EAR-NC^* mutants displayed trichome defects indistinguishable from the null mutant (**Fig. 1D**).

Imaging of primary roots and root hair quantification showed that the wild-type *proGL2:EYFP:GL2* construct rescues the ectopic root hair phenotype of *gl2-5* mutants as described previously (Khosla et al., 2014). In contrast, the N-terminal EAR mutants (*gl2^ΔEAR-N^* and *gl2^EAR-N^*) exhibit a partial root hair patterning phenotype, and the *gl2^EAR-C^* and *gl2^EAR-NC^* mutants display ectopic root hair formation like that of *gl2-5* (**Figs. 2A, 2B**). Both the N-terminal and C-terminal EAR motif mutant transgenes were detected by EYFP expression patterns in the root epidermis, as expected from the GL2 native promoter (**Fig. 2C**). Next, the mutants were examined for seed coat mucilage production (**Fig. 2D**). Similar to phenotypes in leaves and roots, the N-terminal EAR motif mutants displayed a partial phenotype whereas the C-terminal EAR motif mutants appeared similar to the *gl2-5* null mutant. Taken together, we observed that the *gl2^ΔEAR-N^* and *gl2^EAR-N^* mutants exhibit an intermediate phenotype, signifying the importance of the N-terminal EAR motif GL2 activity in specific epidermal cell types including trichomes on leaves, root non-hair cells in the root and seed coat mucilage secretory cells in the seed. In comparison, the C-terminal EAR motif mutants appear to render the GL2 protein completely inactive.

**Figure 2.**
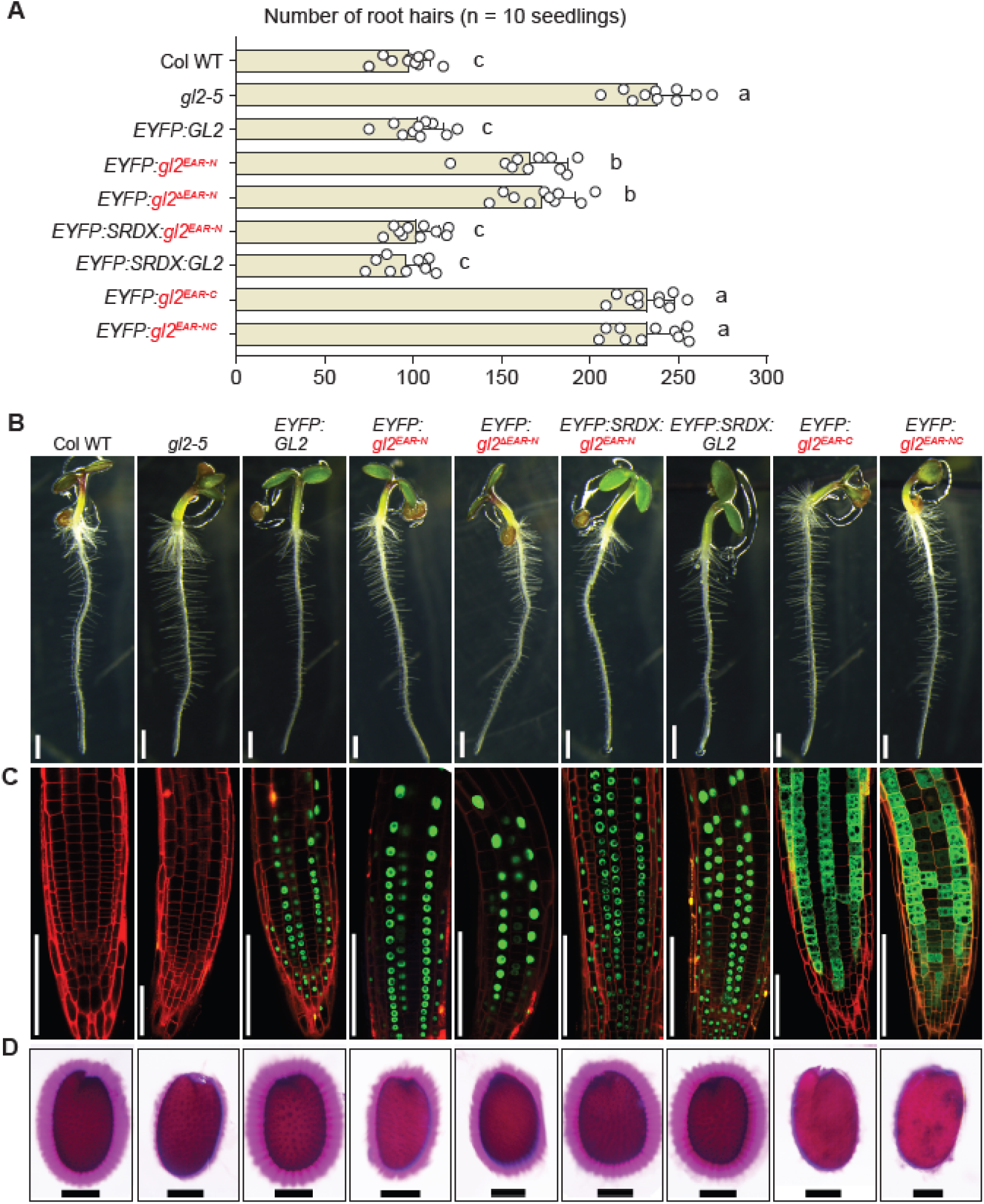
The *gl2* EAR motif mutants display root hair and seed coat mucilage defects. (A) Quantification of root hairs from 4-5 day-old seedlings. Columbia wild type (Col WT) and *EYFP:GL2* transgenic plants exhibit normal root hair formation. In contrast, the *gl2-5* null mutant and the *gl2^EAR-C^* and *gl2^EAR-NC^* EAR-motif mutants display excess root hairs. The *gl2^EAR-N^ and gl2^ΔEAR-N^* N-terminal EAR motif mutants exhibit an intermediate root hair phenotype. Error bars depict the SD, and letters denote significant differences between genotypes for n=10 seedlings as determined by one-way ANOVA and Tukey’s test (p < 0.0001). (B) Representative images of the seedlings used for root hair quantification in **(A)**. Bar = 1 mm. (C) Subcellular localization of EYFP-tagged proteins in primary roots. Col WT and *gl2-5* served as negative controls, lacking EYFP fluorescence. Wild-type GL2 and the *gl2^EAR-N^ and gl2^ΔEAR-N^* N-terminal EAR-motif mutants exhibit nuclear localization, whereas the *gl2^EAR-C^* and *gl2^EAR-NC^* EAR-motif mutants display a defective nuclear localization pattern. Bar = 50 μm. (D) Ruthenium red staining of seeds to visualize seed coat mucilage. Normal seed coat mucilage is observed for Col WT and for *gl2-5* seeds expressing wild-type *EYFP:GL2*, *EYFP:SRDX:gl2^ΔEAR-N^*, and *EYFP:SRDX:GL2*. The *gl2* N-terminal EAR motif mutants display a partial phenotype. In contrast, the *gl2^EAR-C^* and *gl2^EAR-NC^* EAR-motif mutants are defective in seed mucilage production, like *gl2-5*. Bar = 100 µm.

### C-terminal EAR motif mutants show impaired nuclear localization and homodimerization

To investigate the possibility that mutations in the C-terminal EAR motif result in protein misfolding, we monitored the subcellular localization of the EYFP-tagged proteins. While fluorescence was absent in the wild type and *gl2*-5 lines that lacked EYFP-tagged proteins, plants harboring wild-type EYFP:GL2 exhibited nuclear localization, consistent with our previous findings (Ahmad et al., 2024). Similarly, the N-terminal EAR mutant proteins (EYFP:gl2^ΔEAR-N^, EYFP:gl2^EAR-N^) displayed nuclear localization (**Fig. 2C**). In contrast, the EYFP:gl2^EAR-C^ and EYFP:gl2^EAR-NC^ mutant proteins displayed defection subcellular localization, possibly due to protein misfolding and aggregation (**Fig. 2C**).

We investigated the effects of the C-terminal and N-terminal EAR motif mutations on GL2 dimerization activity using yeast Y2H assays (**Fig. 3A**). Western blot analysis of bait proteins indicated comparable expression levels, arguing against the possibility that the nonfunctional proteins are poorly expressed (**Fig. 3B**). Both of the N-terminal EAR-motif mutants demonstrated proficient homodimerization when employed as bait or prey with GL2 or itself, comparable to the GL2 positive control (**Fig. 3A**). In contrast, the gl2^EAR-C^ and gl2^EAR-NC^ mutants failed to form dimers (**Fig. 3A; Fig. S4**). Therefore, we excluded the *gl2^EAR-C^* and *gl2^EAR-NC^* mutants from further analysis, as their defects in nuclear localization and homodimerization likely account for their null mutant phenotypes.

**Figure 3.**
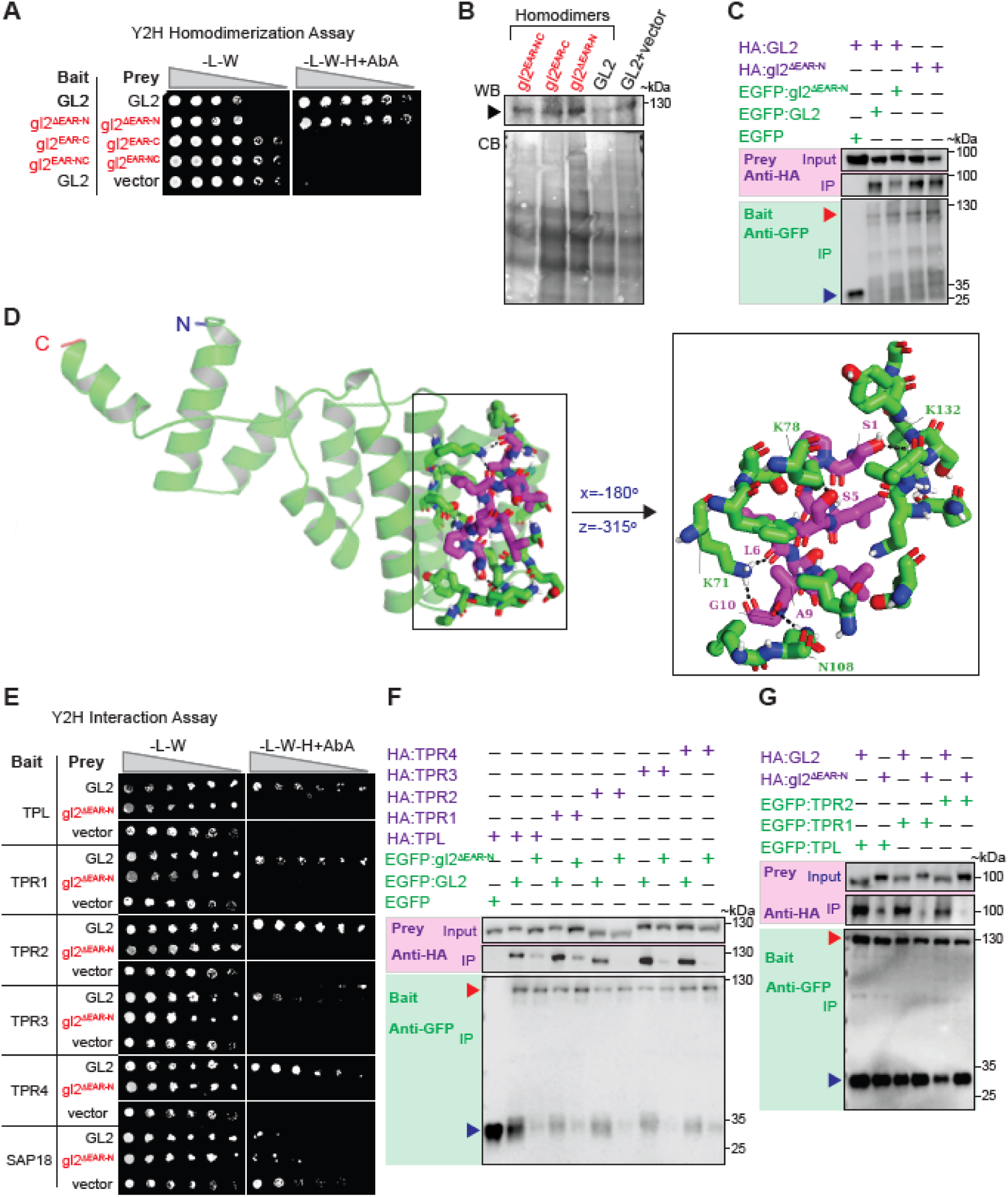
The N-terminal EAR motif is required for GL2 interaction with TPL/TPR corepressors but not for homodimerization. (A) Y2H assays show that gl2^DEAR-N^ mutants exhibit homodimerization similar to that of wild-type GL2, whereas the gl2^EAR-C^ and gl2^EAR-NC^ mutants fail to dimerize. Self-interactions were assayed on selective media lacking Leu, Trp, and His (-L-W-H) in the presence of Aureobasidin A (AbA). GL2 failed to interact with the empty vector control. Four-fold dilutions of yeast are indicated from left to right. (B) Western blotting (WB) with anti-GAL4(DBD) confirmed the expression of the wild-type and mutant bait proteins in the Y2H assays (**A**). Top: Migration of the bait proteins is indicated by the arrowhead. Bottom: Coomassie blue (CB) staining shows protein loading for the WB. (C) Co-IP of proteins expressed in *N. benthamiana* confirms that wild-type GL2 and the gl2^ΔEAR-N^ mutant are proficient in homodimerization. EGFP-and HA-tagged proteins served as baits and preys, respectively. Input and immunoprecipitated (IP) proteins were detected in western blots with anti-GFP or anti-HA antibodies. Predicted kDa sizes are indicated. Red arrowhead marks position EGFP-tagged GL2 proteins; blue arrowhead marks position of EGFP alone or EGFP degradation product. (D) Computational docking of the GL2 EAR motif (SPALSLSLAG) with Arabidopsis TPL. Interacting residues of the EAR motif (magenta) and TPL (green) are labeled. Black dotted lines illustrate hydrogen bonding between the EAR motif and TPL. (E) Y2H assays show that GL2 interacts with TPL and TPR proteins and that the GL2 EAR motif is required for this interaction. The empty vector and SAP18 served as negative controls. Yeast growth on media lacking Leu and Trp (-L-W) signifies presence of bait and prey plasmids, while growth on selective media (-L-W-H) containing AbA indicates protein interaction. Four-fold dilutions are indicated left to right. (F) and **(G)** Co-IP experiments demonstrate GL2-TPL and GL2-TPR interactions using *N. benthamiana* transient expression. EGFP– and HA-tagged proteins served as baits and preys, respectively. Input indicates the initial protein amount used for the Co-IP. Input and immunoprecipitated (IP) proteins were detected in western blots with anti-HA and anti-GFP antibodies. Approximate kDa sizes are indicated. **(F)** Co-IP shows wild-type GL2 in planta interaction with TPL and TPR proteins dependent on the EAR motif whereas EGFP:gl2^ΔEAR-N^ displayed weaker or no interaction. EGFP-tagged GL2 (red arrowhead) or EGFP negative control (blue arrowhead) served as bait whereas HA-tagged TPL or TPR proteins served as preys. **(G)** Reverse Co-IP experiments confirm EAR motif-dependent interactions between TPL/TPR proteins and GL2. EGFP:TPL, EGFP:TPR1 and GFP:TPR2 served as baits, while GL2 and gl2^ΔEAR-N^ served as preys. Red arrowhead marks position EGFP-tagged TPR proteins; blue arrowhead marks position of EGFP alone or EGFP degradation product. TPL, TPR1 and TPR2 exhibit interaction with GL2 but not with gl2^ΔEAR-N^.

The homodimerization proficiency of gl2^ΔEAR-N^ and gl2^EAR-N^ was further validated by transient expression of the proteins in planta, followed by co-immunoprecipitation (co-IP) (**Fig. 3C**). GL2 and gl2^ΔEAR-N^ were reciprocally utilized as bait and prey when tagged with enhanced green fluorescent protein (EGFP) and hemagglutinin (HA) tags, respectively. EGFP as bait, along with GL2 as prey, served as negative control. The results show that gl2^ΔEAR-N^ retained its dimerization activity similar to that of wild-type GL2 (**Fig. 3C**). Overall, the results demonstrate that the N-terminal EAR motif mutants retain homodimerization and nuclear localization proficiency.

### SRDX fusion restores the *gl2^EAR-N^* function

The SRDX domain contains two overlapping EAR motifs (LDLDLELRLGFA), imparting dominant repression activity to associated transcription factors (Hiratsu et al., 2003). We translationally fused *SRDX* to the N-terminus of *gl2^EAR-N^* (**Fig. 1B**) and wild-type *GL2*. The resulting constructs (*proGL2:EYFP:SRDX:GL2* and *proGL2:EYFP:SRDX:gl2^EAR-N^*) were introduced into the *gl2-5* mutant background. Subcellular localization of the corresponding proteins in roots confirmed nuclear expression (**Fig. 2C**). Phenotypic analyses show that *EYFP:SRDX:gl2^EAR-N^* rescued the *gl2-5*, displaying trichomes levels (**Figs. 1C, 1D**), root hair patterning and seed mucilage production similar to wild-type *EYFP:GL2* and Col (**Figs. 2A-2C**). Intriguingly, the fusion of *SRDX* with wild-type *GL2* (*EYFP:SRDX:GL2*) led to slightly fewer trichomes on first leaves, but no detectable differences in root hair and seed coat mucilage formation. Our results demonstrate that the fusion of SRDX with gl2^EAR-N^ restores its function in epidermal development.

### GL2 EAR motif interacts with TPL corepressor in silico

Next we applied molecular docking, a computational approach for protein-protein interaction analysis (Wang et al., 2019), to investigate the putative GL2-derived N-terminal EAR motif interaction with the TPL corepressor, due to availability of a crystal structure (PDB ID: 5NQV). The GL2 EAR motif structure was predicted using PEPFOLD3. Our protein-peptide docking results indicate that the GL2 EAR motif interacts with N-terminal residues of TPL, forming hydrogen bonds and electrostatic interactions (**Fig. 3D; Table S2**). GL2 EAR motif residues Ser5 and Leu6 interact with Lys71 and Lys78 of TPL, consistent with crystal structure of TPL bound to the EAR motif of IAA27 (Martin-Arevalillo et al., 2017). We additionally observed novel hydrogen bonding and electrostatic interactions, possibly due to variations in nearby residues (**Table S2**).

### GL2 EAR motif is required for interaction with TPL and TPR1-4 corepressors in vivo

Given that proteins comprising the EAR motif are predicted to interact with various corepressor proteins (Kagale and Rozwadowski, 2011), we chose TPL, TPR1-4, and SIN3 ASSOCIATED POLYPEPTIDE P18 (SAP18) as potential candidates for interaction with GL2 and gl2^ΔEAR-N^ in Y2H assays. The results indicate that GL2 shows protein-protein interactions with TPL and TPR1-4, while gl2^ΔEAR-N^ fails to interact with any of these corepressors (**Fig. 3E**). TPL and TPR1-4 did not display interactions with empty vectors, indicating specificity of the interactions and the absence of autoactivation activity for these proteins. The expression levels of bait proteins in all combinations were confirmed using western blot analysis (**Figs. S5A, S5B**). In comparison, SAP18 displayed autoactivation activity when employed as bait (**Fig. 3E**). To circumvent this issue, we tested the interaction in reverse, using SAP18 as prey and GL2 and gl2^ΔEAR-N^ as baits. This approach produced similar results, showing that GL2 fails to interact with SAP18 (**Fig. S5C**). Taken together, the findings indicate that GL2 specifically interacts with TPL and TPR1-4, but not SAP18, and that the EAR motif is required for this interaction.

To validate the Y2H data we transiently expressed the proteins in planta followed by Co-IP. The GL2 and gl2^ΔEAR-N^ fusions to EGFP were used as baits, while TPL and TPR1-4 fusions to the HA tag were used preys. Total proteins were extracted, and GFP-Trap Magnetic Agarose beads were used to pull down EGFP:GL2 and EGFP:gl2^ΔEAR-N^, followed by western blotting. Our results show that GL2 coimmunipreciptated TPL and the TPR proteins (TPR1, TPR2, TPR3, and TPR4) (**Fig. 3F**). In contrast, gl2^ΔEAR-N^ failed to interact with the TPL/TPR prey proteins, consistent with our Y2H data. The EGFP control failed to show interaction with HA:TPL, demonstrating specificity. We reciprocally verified these results using TPL, TPR1, and TPR2 as baits, and GL2 and gl2^ΔEAR-N^ as prey, with similar outcomes (**Fig. 3G**). Our results demonstrate that GL2 physically interacts with the TPL/TPR corepressors, whereas gl2^ΔEAR-N^ shows little or no interaction. Taken together, the in planta co-IP results verify our Y2H data, indicating that the GL2 N-terminal EAR motif is required for corepressor interaction.

### Double mutant analysis reveals role of PRC2 components *CLF* and *SWN* in trichome development, likely mediated through GL2-CLF interaction

Previous analysis of TPL1/TPR1-4 multiple mutants illustrates that these genes display functional redundancy and that they are essential for proper development of the epidermis (Long et al., 2006). Therefore, it is challenging to demonstrate their specific role in GL2-mediated trichome formation. PRC2 components are also essential for plant development and promote gene repression by catalyzing trimethylation of histone H3 at lysine 27 (H3K27me3) (Mozgova et al., 2015). *CLF* and *SWN* encode functionally redundant histone methyltransferases that are critical for facilitating epigenetic regulation of gene expression underlying plant growth and development (Shu et al., 2019). Mutants of *PRC2* are implicated in defective trichome development (Chanvivattana et al., 2004; Johnston et al., 2010). Given the functional redundancy between *CLF* and *SWN*, we investigated trichome phenotypes in *clf-1/-;swn-50/+* progeny. Our results show that *clf*;*swn* double mutants exhibit developmental defects, including smaller plant size and reduced trichome numbers with branching defects (**Fig. 4A**). Quantification indicated significant defects in trichome formation and branching in *clf;swn* mutants, with more pronounced effects in homozygotes in comparison to heterozygotes (**Fig. 4B**). The *clf-1/-;swn-50/+* heterozygous plants predominantly produced 2-branched trichomes, with fewer 3– and 4-branched trichomes. In contrast, *clf-1/-;swn-50/-* homozygotes primarily produced 1-branched trichomes, with few 3-branched trichomes and no 4-branched trichomes. Our observations show that *CLF* and *SWN* are critically important for normal trichome development, with *SWN* acting in a dose-dependent manner in the absence of *CLF*.

**Figure 4.**
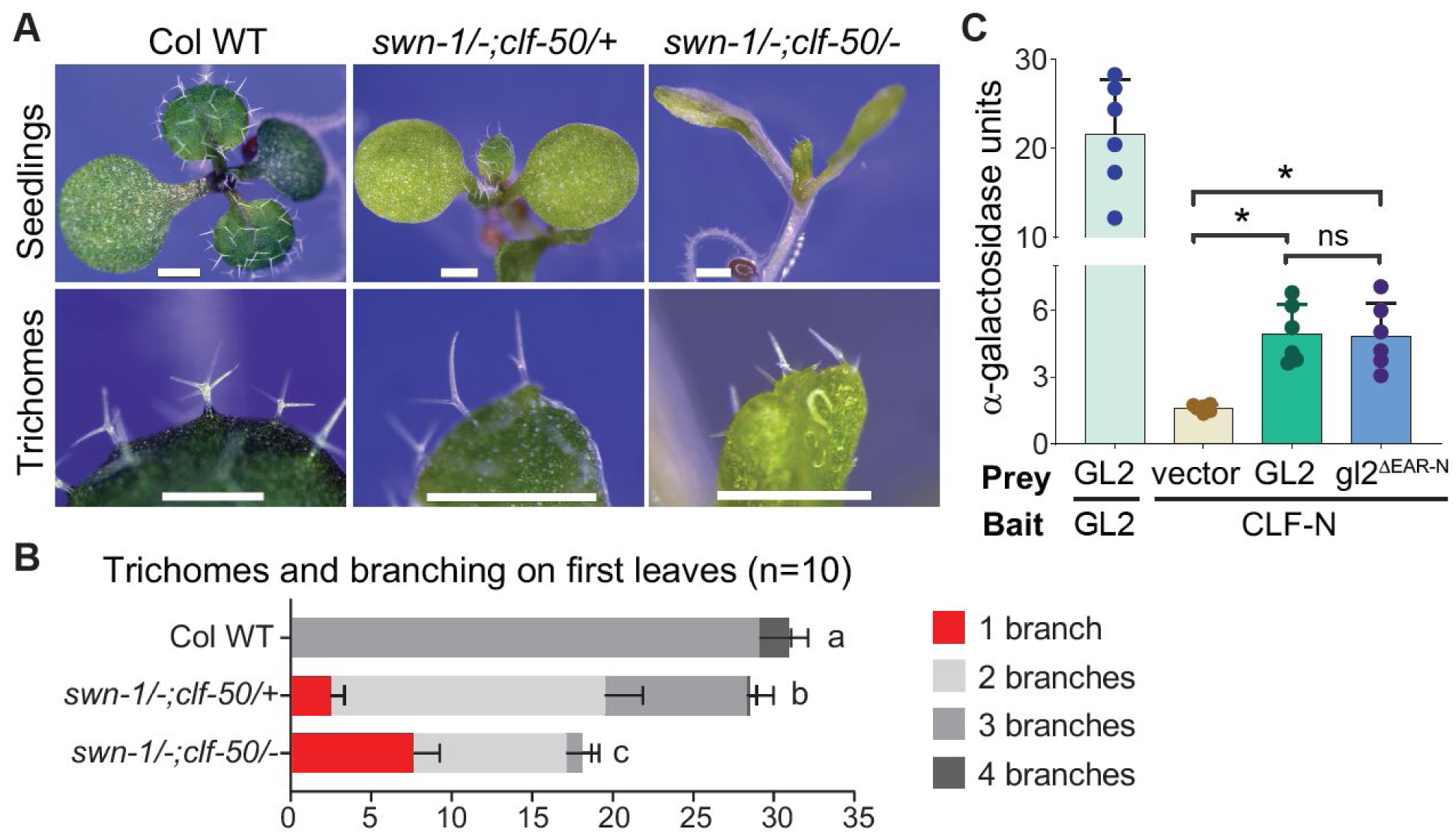
CURLY LEAF (CLF) is critical for trichome development via its direct interaction with GL2. **(A)** Trichome phenotypes on first leaves. Wild-type (WT) plants display normal trichomes having three or more branches. The *swn;clf* double mutants exhibit defective trichomes, and this phenotype is more pronounced in *swn-1/-;clf-50/-* homozygotes versus *swn-1/-;clf-50/+* heterozygotes depending on copy number of *CLF*. Bar = 500 µm **(B)** Quantification of trichome numbers and branching on first leaves. Representative seedlings are shown in **(A)**. Error bars show SD, and letters denote significant differences between genotypes for n=10 seedlings as determined by one-way ANOVA and Tukey’s test (p < 0.001). **(C)** Quantitative α-galactosidase assay indicates interactions between GL2 or gl2^ΔEAR-N^ and CLF-N. The GL2 homodimer served as a positive control; the empty vector in combination with CLF-N served as a negative control. Significant differences for n=6 biological replicates are indicated by asterisk as determined by student’s *t*-test (p < 0.001).

We hypothesized that CLF is critical for trichome formation via its interaction with GL2. A previous study demonstrated that various EAR-motif associated transcription factors recruit PRC2 components to catalyze histone methylation (Baile et al., 2021). We performed Y2H assays and detected an interaction between CLF-N (an N-terminal segment of CLF) and GL2 (**Fig. 4C**). Quantitative analysis demonstrated that GL2 and gl2^ΔEAR-N^ exhibited similar levels of interaction with CLF-N (**Fig. 4C**). These unexpected results suggest that the GL2 N-terminal EAR motif is not required for the interaction with CLF-N and involves alternative mechanism.

### Transcriptome analysis highlights the role of N-terminal EAR motif for proper GL2 activity in regulating gene expression

To investigate how the GL2 N-terminal EAR motif contributes to transcriptional regulation, we performed RNA-seq on seedling roots from wild type, the *gl2-5* null mutant, and the EAR-motif mutant *gl2^ΔEAR-N^* (**Fig. 5; Datasets S1-S6**). Principal component analysis (PCA) revealed a clear separation of genotypes, with wild type and *gl2-5* at opposite extremes, and *gl2^ΔEAR-N^* positioned between them, indicating an intermediate global transcriptomic state (**Figure S6**). Hierarchical clustering and heatmaps of the differentially expressed genes (DEGs) (padj<0.05, log2FC>1) further supported this pattern and confirmed high reproducibility among biological replicates.

**Figure 5.**
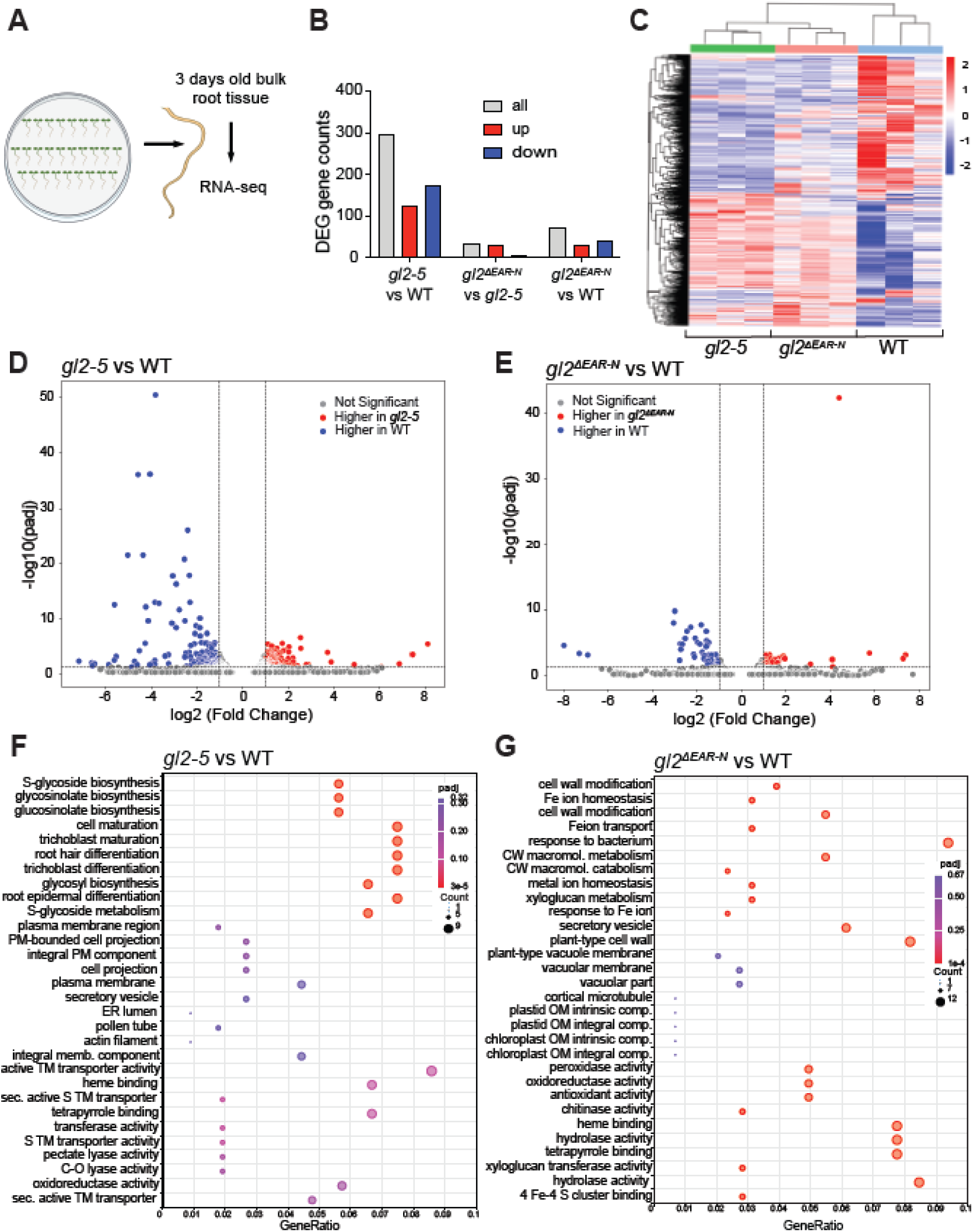
*gl2^ΔEAR^* shows intermediate global gene expression patterns. **(A)** Pooled root tissue from 3-day-old Arabidopsis seedlings used for RNA-seq analysis. **(B)** Differentially expressed gene (DEG) counts in *gl2-5* vs wild type (WT), *gl2^ΔEAR^* vs. *gl2-5*, and *gl2^ΔEAR^* vs. WT. Genes were called differentially expressed with log₂ fold change ≥ 1 and adjusted P value < 0.05 (DESeq2). Total gene counts, up-regulated gene counts (positive log fold change, LFC), and down-regulated gene counts (negative LFC) are indicated. **(C)** Hierarchical clustering heatmap of the union of DEGs from all pairwise comparisons (*gl2-5* vs WT, gl2^ΔEAR^ vs WT, *gl2^ΔEAR^* vs *gl2-5*). Expression values were first transformed as log₂(FPKM+1) and hierarchically clustered; for display, each gene (row) was standardized to a Z-score. The color scale (−2 to 2) reflects row-normalized expression (red = higher, blue = lower relative to the mean). Columns represent biological replicates. **(D and E)** Volcano plots of differential expression for *gl2-5* vs. WT **(D)** and *gl2^ΔEAR^* vs. WT **(E)**. The x-axis in each plot shows log₂ fold change and the y-axis shows log₁₀ (adjusted p value, padj). Red points are higher in *gl2-5* **(D)** or *gl2^ΔEAR^* **(E)**. Blue points are higher in WT. Vertical dashed lines mark log₂FC = 1, and the horizontal dashed line marks padj = 0.05 (DEG threshold). **(F and G)** The x-axis in each plot shows Gene Ratio (query genes annotated to a term / total query genes); point size indicates the number of genes in each term, and point color reflects padj. Terms from Biological Process, Molecular Function, and Cellular Component are shown. Enrichment was assessed using an over-representation analysis (hypergeometric test) with the Benjamini–Hochberg correction against all tested genes. **(F)** Plot showing the gene ontology (GO) enrichment of the genes upregulated in *gl2* compared to WT shown in **(D)**. **(G)** Plot showing the gene ontology (GO) enrichment of the genes downregulated in *gl2^ΔEAR^* compared to WT shown in **(E)**.

In the *gl2-5* comparison to wild type, we identified 297 differentially DEGs (125 up– vs. 172 downregulated), reflecting the major regulatory role of GL2 in epidermal differentiation and cell wall associated processes. Comparison of *gl2^ΔEAR-N^* with wild type identified 70 DEGs (29 up– vs. 41 downregulated), considerably fewer than in the *gl2* and wild type comparison. This reduction in DEGs supports our hypothesis that *gl2^ΔEAR-N^* displays a partially rescued transcriptome, consistent with its intermediate phenotype. Notably, the proportion of upregulated genes (∼40%) is approximately the same in both comparisons. In contrast, the *gl2^ΔEAR-N^* and *gl2-5* comparison identified 32 DEGs (28 up– vs. 4 downregulated) (88% upregulated), indicating the intermediate phenotype of *gl2^ΔEAR-N^* may primarily be explained by a regulatory mechanism that does not require the EAR motif.

Genes that were found to exhibit an intermediate gene expression pattern (down-regulated in comparison to wild-type; upregulated in comparison to *gl2-5*) could contribute to the intermediate root hair phenotype of *gl2^ΔEAR-N^*. We considered such genes to be GL2 dependent but EAR-motif independent. These include cell wall metabolism-related genes (*XTH33/At1g10550*, *EXPA12/At3g15370*), and cell wall modification genes (*XTH17/At1g65310*, *XTH18/At4g30280*, *XTH30/At1g32170*, *XTH33/At1g10550*). Others include a cuticular wax biosynthesis gene *CER4/At2g43590*, and the putative galactose oxidase gene *RUBY/At1g19900* which affects pectin formation. Additionally, some fatty acid related genes (*KSC5/At1g25450*, *ADS1.4/At1g06120*, *CRSH*/*At4g17470*) exhibit an intermediate gene expression pattern. A few genes involved in root morphogenesis and lateral root development (*AIR1;At4g12550*, *EXPA17;At4g01630*) also displayed an intermediate transcript level.

We defined EAR motif-dependent DEGs as those that were similarly expressed in *gl2-5* and *gl2^ΔEAR-N^*. Several of these genes (*IMA4/At1g07367*, *IMA6/At1g07373*) encode long non-coding RNAs related to the regulation of metal ion transport. The gene *CLH1/AT1G19670* involved in the chlorophyll catabolic process and *DIN11/At3g49620*, which is involved in the cellular response to starvation, are upregulated in both *gl2-5* and *gl2^ΔEAR-N^*, suggesting that their expression pattern is EAR-motif dependent. Strikingly, genes involved in glucosinolate metabolic process (BCAT4/At3G19710), IMS2 (At5G23020), ATSDI (At5G48850), SDI2 (At1G04770), MAM1 (At5G23010), and FMO_GS-OX2 (At1G62540)) are broadly upregulated in *gl2-5* compared to the wild type, but are not upregulated in the EAR motif mutant. An exception to this trend was *BGLU32*, encoding beta-glucosidase, which remained upregulated in *gl2^ΔEAR-N^* relative to wild type, suggesting incomplete rescue of glucosinolate breakdown in the partial phenotype mutant.

Very few genes were uniquely downregulated in *gl2^ΔEAR-N^* compared to *gl2*. These included the *QQS/At3g30720* and *SWEET3/At5g53190* genes that are involved in regulating starch metabolic processes and sugar transporter activity. Taken together, the RNA-seq data reveal that GL2 regulates distinct transcriptional modules through both EAR-dependent and EAR-independent developmental pathways, and that the *gl2^ΔEAR-N^* mutant displays a partially rescued transcriptome, consistent with its intermediate phenotype.

## Discussion

### N-terminal EAR motif is conserved in GL2 orthologs

The focus of this work was to study the gene repression mechanism of GL2, a well-studied member of the HD-Zip IV family and a crucial determinant of cell fate in various epidermal cell types. We demonstrate that GL2 harbors an N-terminal EAR motif that is required for the recruitment of TPL/TPR corepressors, thus facilitating gene repression. Sequence analyses suggested that GL2 possesses two putative EAR motifs, the first at the N-terminus upstream of the HD, and the other at the C-terminus (**Fig. 1A**). It is well established that some transcriptional regulators contain more than one EAR motif (Chow et al., 2023). For example, INDOLE-3-ACETIC ACID 7 (IAA7), which participates in auxin-mediated processes in *Arabidopsis*, comprises two EAR motifs, with one being of minor importance compared to the other (Lee et al., 2016). The N-terminal EAR motif of GL2 is highly conserved among its orthologs across the monocots, including maize, rice, and wheat, as well as the dicots, such as cotton, soybean, and tomato (**Fig. S3**). SpCH4, an ancestral HD-Zip IV in the charophycean green algae *Spirogyra pratensis,* contains an N-terminal EAR motif similar to that of GL2 (**Table S1**). This conservation indicates the functional significance and evolutionary retention of this motif. In comparison, the C-terminal EAR motif of GL2 was found to be less conserved. Notably, PROTODERMAL FACTOR2 (PDF2), the closest homolog of SpCH4, and its paralog, *Arabidopsis thaliana* MERISTEM LAYER1 (ATML1), lack an LxLxL-type EAR motif (**Table S1**). The absence of an EAR motif in a subset of HD-Zip IV proteins points to evolutionary diversification in their mechanisms of gene regulation.

### Activation versus repression roles of GL2

Our RNA-seq data suggest that *GL2* has both EAR motif-dependent and EAR motif-independent regulatory effects. Comparison of DEG patterns showed that deletion of the EAR motif partially restores the transcriptional defects of the *gl2-5* null mutant. Genes involved in cell-wall modification, cuticle and lipid metabolism, fatty acid biosynthesis, and root morphogenesis were strongly downregulated in *gl2-5* but showed intermediate expression levels in *gl2^ΔEAR-N^*, indicating EAR motif-independent GL2 activation functions. A substantial number glucosinolate biosynthetic pathway genes were upregulated in *gl2-5* but not in *gl2^ΔEAR-N^*, potentially due secondary effects from the absence of GL2 activity. A small group of genes, including *QQS* and *SWEET3*, was uniquely downregulated in *gl2^ΔEAR-N^* but not in *gl2-5*, indicating additional transcriptional consequences specific to EAR motif removal.

Various lines of evidence suggest that GL2 requires balanced transcriptional activation and repression activities to regulate normal epidermal differentiation. Conversion of GL2 into a transcriptional activator by N-terminal-fusion with VP16 resulted in ectopic root hairs in roots as well root hair-like structures in the hypocotyl epidermis (Ohashi et al., 2003). A previous report suggested that GL2 physically interacts with the adaptor proteins GIR1 and GIR2 to facilitate TPL-dependent transcriptional repression of GL2 targets (Wu and Citovsky, 2017a, b). Here we demonstrate that GL2 does not require an adaptor protein since it itself contains a functional EAR motif at its N-terminus. Our mutational analysis showed that this N-terminal EAR motif is critical for GL2 function (**Figs. 1C, 1D**; **Figs. 2A, 2B**). However, the epidermal defects in the N-terminal EAR mutant are not as severe as those of the *gl2-5* null mutant. It is unlikely that C-terminal EAR motif plays a minor role in corepressor binding since our experiments show that the gl2^ΔEAR-N^ mutant fails to interact with the TPL/TPR proteins (**Fig. 3D**). Another possibility is that the gl2^ΔEAR-N^ mutant unlike the null mutant, retains its activation activity, and the observed phenotype is the result of positive target gene expression. We generated substitution mutants to explore the function of the putative C-terminal EAR motif. Unfortunately we were unable to establish the role of this C-terminal EAR motif since the corresponding mutant proteins exhibited nuclear localization defects (**Fig. 2C**) and a null mutant phenotype (**Figs 1C, 1D**; **Figs 2A, 2B**), possibly due to issues in protein folding.

BRASSINAZOLE RESISTANT 1 (BZR1) is an example of an EAR-motif containing transcription factor that regulates both gene activation and repression, similar to GL2 (He et al., 2005; Sun et al., 2010; Oh et al., 2014). BZR1 EAR motif deletion results in the loss of activation and repression activities, and SRDX fusion rescues both, suggesting a more complex role for the EAR motif in gene regulation (Oh et al., 2014). Similarly, our transcriptome data for the GL2 N-terminal EAR motif mutant uncovers changes in both positive and negative gene expression. Identifying an activation domain in GL2 and investigating its interplay with the EAR motif will provide a better understanding of the plasticity of GL2-mediated transcriptional activities

Our results demonstrate that the N-terminal fusion of SRDX with *gl2^EAR-N^* restores its activity and results in normal epidermal development. Thus, the GL2 EAR motif that is present at its N-terminus serves a similar repressive function as SRDX. Intriguingly, SRDX fusion with wild-type GL2 led to a decrease in trichome formation (**Fig. 1D**), suggesting that the additional EAR motif in SRDX interferes with GL2 function. The presence of multiple EAR motifs could facilitate the simultaneous binding of multiple corepressors in addition to TPL/TPR proteins. A previous study found that SAP18 interacts with SlEAD1, which contains two overlapping EAR motifs with sequence similarity to SRDX (Wang et al., 2020a). Root hair quantification showed no significant differences between *EYFP:GL2* and *EYFP:SRDX:GL2* lines, possibly because of distinct regulatory mechanisms in the different tissues (**Fig. 2A**).

### EAR motif is required for interaction between GL2 and TPL or TPR corepressors

The TPL and TPR proteins contain an N-terminal TOPLESS domain (TPD) that is required for interaction with EAR motif-containing proteins (Szemenyei et al., 2008; Plant et al., 2021). Crystal structure analysis of *Arabidopsis* TPL TPD (180 residues) in complex with the EAR motif (TELRLGLPG) of auxin/indole-3-acetic acid (IAA27) protein reveals that they associate via hydrophobic residues present in the TPD (Martin-Arevalillo et al., 2017). The interacting residues exhibit high conservation among the five Arabidopsis TPL/TPR proteins and their orthologs across the plants (Martin-Arevalillo et al., 2017). Therefore it is not surprising that the *TPL/TPR* genes redundantly determine apical embryonic fate, and a quintuple mutant *tpl-2; tpr1-1; tpr3-1; tpr4-1*; TPR2-RNAi displays a similar phenotype as *tpl-1* (Long et al., 2006). Our interaction data demonstrates that GL2 directly interacts with TPL and TPR1-4 through its N-terminal EAR motif (**Figs. 3E-3G**). While GL2 interacts with the TPL/TPR corepressors, it appears not to interact with SAP18 (**Fig. 3E**). This mirrors the interaction profile of TIE1, an EAR motif-containing transcriptional corepressor that regulates leaf development (Tao et al., 2013).

Co-expression analysis indicates that GL2, TPL, and TPR proteins are variably co-expressed in the trichomes, roots, and the seed coat (**Fig. S6**). This co-expression pattern, combined with our protein interaction data, suggests that TPL/TPR corepressors function redundantly in regulating GL2 activity. However, the mechanism by which GL2 selects between the TPL and TPR isoforms remains unclear. The factors potentially influencing this selection include, but are not limited to, post-translational modifications, expression levels or availability of TPL and TPR proteins, and competition from other EAR-motif containing proteins.

### Regulation of EAR motif mediated gene repression

While our study offers initial insights into the EAR motif-mediated gene repression mechanism of GL2, post-translational modifications, such as phosphorylation, may positively or negatively impact EAR motif-mediated gene repression by modulating protein-protein interactions, subcellular localization, and/or turnover (Kagale and Rozwadowski, 2011). Ser and Thr residues associated with the EAR motif are found to be phosphorylated in several proteins, such as INDOLE-3-ACETIC ACID INDUCIBLE 9 (IAA9), ETHYLENE RESPONSE FACTOR 10 (ERF10), and BES1/BZR1 HOMOLOG 4 (BEH4) (Kagale et al., 2010). Intriguingly, GL2 is Ser-rich and contains five Ser and one Thr residue upstream of the N-terminal EAR motif. Additionally, the GL2 N-terminal EAR motif (LSLSL) contains two Ser residues (Ser23 and Ser25), with Ser23 being highly conserved among GL2 monocot and dicot orthologs (**Fig. S3**). Future studies investigating whether phosphorylation affects the association of GL2 with corepressor proteins could provide valuable insights into the mechanisms that enable GL2 to function as either a transcriptional activator or repressor in a spatiotemporally regulated manner.

### GL2 interacts with CLF independently of the N-terminal EAR motif

A previous study showed that the EAR motif is sufficient for interaction and recruitment of chromatin remodeling proteins including histone methyltransferases and histone deacetylases (Baile et al., 2021). Our findings show that the GL2 N-terminal EAR motif is not required for interaction with the CLF histone methyltransferase (**Fig. 4C**). It is possible that the GL2 C-terminal EAR motif facilitates the interaction with CLF. Trichome quantification in *swn/-;clf/+* and *swn/-;clf/-* mutants showed branching defects (**Figs. 4A, 4B**), further supporting its role in GL2-mediated trichome cell-type differentiation. The CLF and SWN act redundantly (Shu et al., 2019), but we failed to detect interactions between SWN and GL2 in Y2H assays, suggesting an alternative mechanism for its recruitment.

Overall, our study shows that GL2 orchestrates gene expression of its transcriptional targets through its association with TPL/TPR corepressors and the CLF histone methyltransferase (**Fig. 6**). Our future work will investigate how these associations are themselves regulated and whether these two groups of proteins act synergistically or in a context-dependent manner. Elucidating the interplay between GL2 and its corepressors and associated chromatin-remodeling enzymes will provide deeper mechanistic insight into the complex gene regulation underlying epidermal development.

**Figure 6.**
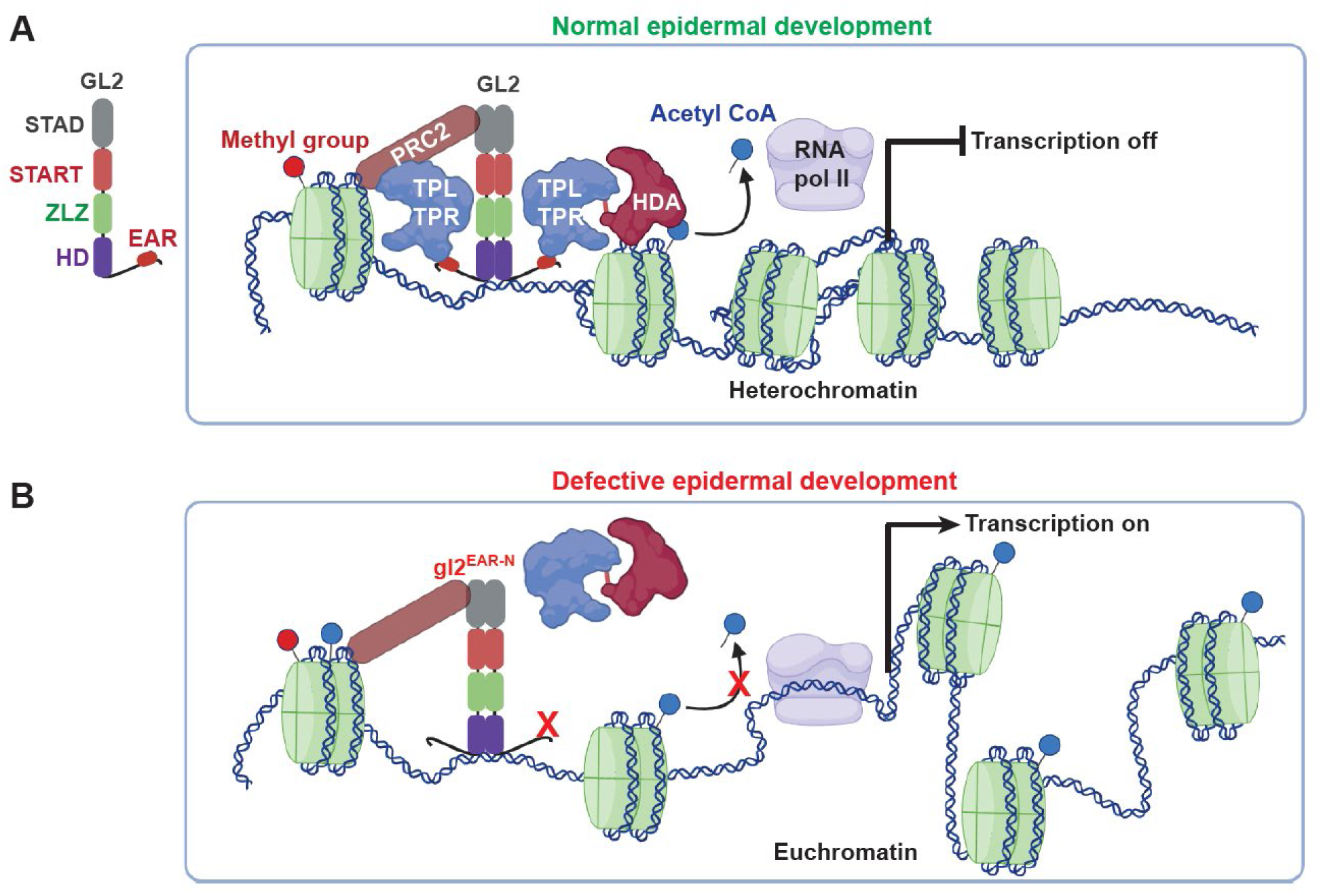
Model for GL2-mediated gene repression in controlling epidermal development. **(A)** In wild type, GL2 binds as a dimer to promoters of its transcriptional targets to facilitate the recruitment of TPL/TPR corepressor proteins via its EAR motif. In turn, corepressor proteins recruit chromatin remodeling proteins. GL2 additionally associates with PRC2 components through corepressors and/or direct interactions, such as with CLF, independent of the EAR motif, resulting in chromatin marks for repression of target genes. **(B)** In gl2^ΔEAR-N^, the absence of the EAR motif results in the loss of interaction between GL2 and corepressor proteins and, ultimately, the loss of HDAC recruitment. However, gl2^ΔEAR-N^ retains its dimerization activity and interaction with CLF, leading to histone methylation associated with partial repression of transcriptional targets, consequently resulting in defective epidermal development and partial *gl2* mutant phenotypes.

## Materials and methods

### Plant materials and growth conditions

Arabidopsis plants utilized in this study were of the Columbia (Col) ecotype, unless otherwise specified. The *gl2-5* null mutant and *proGL2:EYFP:GL2* transgenic lines were previously described (Khosla et al., 2014). The *swn-1/-;clf-50/+* (Wassilewskija (Ws) ecotype) seeds were kindly provided by Justin Goodrich and are described in (Chanvivattana et al., 2004). Seeds were sown on BM6 soil supplemented with vermiculite and perlite at a ratio of 4:2:1. After stratification at 4°C for 4-5 days, the plants were grown at 23°C and continuous light. The *Agrobacterium tumefaciens* GV3101 (pMP9) floral dip transformation method (Clough and Bent, 1998) was used to construct transgenic lines in the *gl2-5* genetic background. Multiple transformants were selected on 20 mg/ml Hygromycin B prior to transferring to soil. T1 plants were screened based on enhanced yellow fluorescent protein (EYFP) expression and segregation according to Mendelian 3:1 ratios on Hygromycin B. For each construct, 2-3 independent transformants were analyzed, and at least one representative line was genotyped by sequencing and taken for further characterization.

### Construction of plasmids and binary vectors

EAR motif Ala substitution (*gl2^EAR-N^* and *gl2^EAR-C^*) and deletion (*gl2^ΔEAR-N^*) mutants were constructed with the Q5 Site-Directed Mutagenesis Kit (E0554S, New England Biolabs, Ipswich, MA, USA). This process used the pENTR/D-TOPO vector containing wild-type *GL2* cDNA as a template. The *gl2^EAR-NC^* mutant was created using the pENTR/D-TOPO vector harboring *gl2^EAR-C^* cDNA. The mutants were validated by sequencing and cloned into a previously described binary vector SR54 (p*roGL2:EYFP:GL2*) (Khosla et al., 2014). The inserts were amplified with Q5 High Fidelity DNA polymerase (M0491, New England Biolabs). Amplicons were purified using Nucleospin Gel and PCR Clean-up kit (Macherey-Nagel). The SR54 vector was first linearized using the *Sal*I and *Kpn*I restriction enzymes, and amplicons were assembled into the linearized vector using the NEBuilder HiFi DNA Assembly kit (New England Biolabs). The *EYFP:SRDX:gl2^EAR-N^* and *EYFP-SRDX-GL2* constructs were generated by *Sal*I digestion of the SR54 vector containing *proGL2:EYFP:gl2^EAR-N^* and *proGL2:EYFP:GL2*, followed by assembly with the *SRDX* oligonucleotide using the NEBuilder HiFi DNA Assembly mix. Primers and other oligonucleotides used for cloning are listed in **Table S3**.

To construct the Y2H plasmids, the cDNAs of full-length *GL2*, *gl2^ΔEAR-N^*, *gl2^EAR-C^*, and *gl2^EAR-NC^* were transferred to the pDEST22 and pDEST32 vectors from pENTR/D-TOPO plasmids. TPR2-4 were transferred from the entry vector pDONR207, kindly provided by Barry Causier, to pDEST32, using the Gateway LR Clonase II Enzyme mix (ThermoFisher Scientific). TPL and TPR1 was cloned into the pENTR/D-TOPO vector using a directional pENTR/D-TOPO Cloning Kit (Invitrogen, Carlsbad, CA, USA). PRC2 genes (*EMF2-C*, *CLF-N*, and *FIE*) cloned into pENTR/D-TOPO and *pDEST32-MSI1* were gifts from Doris Wagner and are described in (Xiao et al., 2017). *pENTR223-SWN* was acquired from the Arabidopsis Biological Resource Center. Genes cloned in entry vectors were subsequently transferred to the pDEST32 vector via Gateway cloning.

Binary vectors used for Co-IP were generated by Gateway cloning as above, transferring GL2 wild-type and mutant cDNA sequences to the C-terminus of the enhanced green fluorescent protein (EGFP) tag in the pK7WGF2 or hemagglutinin (3xHA) tag in the pEarleyGate 201 destination vectors, with pENTR/D-TOPO serving as the donor vector. TPL and TPR1 were transferred from pENTR/D-TOPO, whereas TPR2, TPR3, and TPR4 were transferred from pDONR207 to pEarleyGate 201 or pK7WGF2.

### Phenotypic characterization and microscopy

To image seedlings, trichomes, primary roots, and seeds, we used a Leica M125 fluorescence stereo microscope equipped with a GFP2 filter and a Leica DFC295 digital camera. Trichome quantification of was performed as previously described (Khosla et al., 2014). The *swn-1/-;clf-50/+* progeny and wild-type Columbia seedlings were grown on 1X Murashige and Skoog (MS) media containing 0.8% plant agar and 1% sucrose to quantify trichome numbers and branching patterns. To evaluate and quantify root hairs, sterilized seeds were plated on 1X MS supplemented with 0.4% phytagel [*w/v*] (P8169, Sigma Aldrich) and subjected to stratification at 4°C for 3-4 days. The seedlings were grown vertically at 23°C under continuous light for 3 days. Seed coat mucilage was visualized by ruthenium red (R2751; Sigma-Aldrich) staining, as previously described (Ahmad et al., 2024). Approximately 25-30 seeds per sample were placed in a 24-well plate followed by the addition of 800 µL of 50 mM EDTA and incubation for 2 h. EDTA was removed, and seeds were stained for 4 h with 800 µL of 0.01% ruthenium red [*w/v*]. The stain was removed, and the seeds were rinsed with dH_2_O [pH 7.5] prior to imaging.

Subcellular localization of EYFP-tagged proteins in primary roots of 4-5-day-old seedlings was examined using a Zeiss LSM 700 microscope. Following vapor sterilization, seeds were sown on 0.8% agar plates and stratified at 4°C for a minimum of three days before growing vertically at 23°C under continuous light for 4-5 days. Cell boundaries were stained by treating seedlings with 10 μg/ml propidium iodide (PI) [*w/v*] for 2 min. Excess stain was removed by washing, and the seedlings were subsequently mounted in water on a glass slide. EYFP was excited using a 488 nm laser, with the absorbance spectrum set at 505-600 nm. Concurrently, PI was excited at 555 nm and detected at 610-800 nm. Images were acquired using a 40X oil immersion objective, and processing was performed with Zeiss ZEN Microscopy software.

### GL2 EAR motif and TPL protein-protein docking

In-silico protein-protein interactions between GL2 (EAR motif) and TPL were investigated using a multistep computational approach. The EAR motif (SPALSLSLAG) 3D structure was predicted using PEP-FOLD3 (Lamiable et al., 2016). The TPL 3D structure, comprising N-terminal 184 residues, was retrieved from PDB (PDB ID: 5NQV). Using the UCSF Chimera (Eric F. Pettersen, 2004), we removed the already bound EAR peptide and additional chains from the TPL structure, retaining only chain A. Energy minimization was performed to reduce entropy, and hydrogen atoms were added. The High Ambiguity Driven protein-protein Docking (HADDOCK) web server (De Vries et al., 2010) was used to dock the 3D structures of GL2 EAR and TPL. PyMOL was used to analyze the docked complexes and render images.

### Y2H assay and α-galactosidase activity quantification

Bait (pDEST32) and prey (pDEST22) plasmids were transformed into Y2HGold and Y187 opposite mating-type yeast strains using the standard LiAc/PEG method (Gietz and Woods, 2002). Transformants were selected for the bait and prey plasmids on selective media lacking Leu (-L) and Trp (-W), respectively. The strains were mated on Yeast Peptone Dextrose Adenine (YPDA) medium, followed by growth on –L-W medium for diploid selection. Diploids were grown in liquid media to an OD_600_ of 1.0. Four-fold serial dilutions were prepared from the normalized cultures and placed on permissive and selective media using a 48-pin multiplex plating tool (“Frogger,” Dankar, Inc.), followed by incubation at 30°C for 4-6 days and imaging. Expression levels of baits were confirmed by extracting the total yeast proteins using the sodium hydroxide/trichloroacetic acid method, followed by western blot with the GAL4 [DBD] (RK5C1) mouse monoclonal antibody (sc-510 HRP, Santa Cruz Biotechnology).

The interaction of CLF-N with GL2 and gl2^ΔEAR-N^ was quantified from α-galactosidase units from the *MET1* reporter. Six independent transformants were analyzed for each interaction pair, with three technical replicates per transformant. Diploid yeast were cultured in –L-W liquid media at 30°C. After recording the OD_600_, cells were pelleted and resuspended in 100 μL Z-buffer, containing 50 mM β-mercaptoethanol. Cells were lysed by three freeze-thaw cycles, followed by centrifugation to obtain the cell lysates. For each reaction, 30 μL of the lysate was transferred to a 96-well plate, and 80 μL of 4-nitrophenyl-α-D-galactopyranoside (10 mg/mL) (CAS 7493-95-0) was added. Lysates were incubated at 30°C followed by addition of 190 μL 1 M Na_2_CO_3_. The OD_600_ of the cell culture (before lysis) and the OD_405_ of the reactions were measured using a Bio-Tek Epoch microplate spectrophotometer (Agilent). Calculations were performed according to the following formula: α-Galactosidase units = (Vr × OD_405_)/(t × Vc × OD_600_), where Vr = volume of cleared cell lysate (μL); t = time elapsed (min); Vc = inital cell culture volume (mL)

### Co-IP assay

For protein transient expression and co-immunoprecipitation, the binary vectors pK7WGF2, pEarleyGate 201, and p19 were independently transformed into Agrobacterium GV3101. Cultures were grown overnight until the OD_600_ reached 1.0. Cultures with individual plasmids were combined in a 1:1:0.5 ratio, pelleted, and resuspended in infiltration medium (10 mM MES [pH 5.6], 10 mM MgCl2, 150 mM acetosyringone, and glucose [6 mg/ml]), followed by incubation for 1 h at room temperature in the dark. Leaves from five-week-old *Nicotiana benthamiana* plants grown at 23°C under a 14 h light/10 h dark regime were infiltrated with the resuspended cells using a 1 ml needleless syringe. Plants were moved to the growth chamber for an additional 48 h and analyzed for EGFP expression using a Leica M125 fluorescence stereo microscope equipped with a GFP2 filter. Expressing parts of the leaves (∼0.5 g) were flash frozen and stored at –80°C. For protein extraction, 0.5 g of leaf material co-expressing bait and prey proteins was ground in liquid nitrogen and transferred to 2.5 ml extraction buffer (150 mM NaCl, 50 mM Tris-HCl [pH 7.5], 5% [v/v] glycerol, 1 mM EDTA, 0.5% [v/v] Triton X-100, 1 mM dithiothreitol (DTT), 1X protease inhibitor cocktail (P9599, Sigma Aldrich), 1 mM phenylmethanesulfonylfluoride (PMSF), 2% [*w/v*] polyvinylpolypyrrolidone (PVPP)). The lysate was centrifuged thrice at 20,000 *× g* for 10 min at 4°C to remove cell debris. Cleared lysate was incubated with ChromoTek GFP-Trap Magnetic Agarose (Proteintech) for 1 h at 4°C with continuous end-to-end rotation, followed by centrifugation at 1500 *x g* for 2 min. The supernatant was discarded, and the beads were washed with 1 ml of wash buffer (150 mM NaCl, 50 mM Tris buffer [pH 7.5], 5% [v/v] glycerol, 1 mM EDTA, 0.5% [v/v] Triton X-100, 1 mM DTT, 1X protease inhibitor cocktail (P9599, Sigma Aldrich),1 mM PMSF) at least five times, with each wash lasting for 5 min with continuous rotation. The beads were boiled in Laemmli buffer at 95°C for 10 min and stored at –20°C for western blot (WB) analysis. Lysates were heated with equal amount of 2X Laemmli buffer at 95°C for 10 min and used as the input control, followed by SDS-PAGE and WB using mouse anti-GFP (11814460001, Roche; 1:2000) and rat anti-HA (11867431001, Roche; 1:5000) primary antibodies, and goat anti-mouse and anti-rat (AP136P, Millipore; 1:3000) secondary antibodies. Proteins were detected with SuperSignal West Femto Maximum Sensitivity Substrate (ThermoScientific) using an Azure c300 chemiluminescence imager (Azure Biosystems).

### RNA extraction and RNA sequencing

Sterilized seeds from each genotype were sown on 0.5X MS media, and stratified at 4°C for 6 days. Plates were placed in the 22°C growth chamber vertically for 3 days, followed by harvesting of root tissue samples in liquid nitrogen and brief storage at –80°C. RNA extraction was performed using the QIAGEN RNeasy Plant Mini Kit (74904) and the RNase-free DNase Set (79254). RNA purity and concentration was determined with the Agilent 2100 Bioanalyzer. Three biological replicates from each genotype were selected for cDNA library construction, Illumina sequencing, and bioinformatic analysis by Novogene (Sacramento, CA). Genes were classified as differentially expressed if their adjusted p value ≤ 0.05 and log2(fold change) ≥ 1. Mainstream hierarchical clustering grouped genes by their FPKM values. Gene ontology and KEGG enrichment analyses were performed for all genes with an adjusted p value < 0.05.

## Statistical analysis

GraphPad Prism 7 software was used to perform standard statistical analyses. Comparisons between two groups were evaluated using an unpaired *t*-test with Welch’s correction. One-way ANOVA and Tukey’s multiple comparison tests were used for more than two groups. Statistical significance of the differences is indicated by letters or asterisks as described in the figure legends.

## Accession numbers

GL2, At1g79840; TPL, At1g15750; TPR1, At1g80490; TPR2, At3g16830; TPR3, At5g27030; TPR4, At3g15880; SAP18, At2g45640; CLF; At2g23380; SWN, At4g02020

## Acknowledgments

We thank Barry Causier, Justin Goodrich, Doris Wagner and the ABRC for providing biological materials.

## Author contributions

K.S. and B.A. designed research; B.A., A.U., A.K.B., L.R.M. and K.S. performed research; B.A., A.U, A.K.B., L.R.M. and K.S. analyzed data; and B.A., A.U. and K.S. wrote the paper.

## Data availability

The RNA-seq data generated in this study is available at the NCBI Gene Expression Omnibus under accession number GSE344912.

**The following Supporting Information is available for this article**:

**Figure S1.** Suppression of *gl2* trichome defects by *gir1* mutation.

**Figure S2.** GL2 fails to interact with GIR1 and GIR2 in yeast two-hybrid (Y2H) assays.

**Figure S3.** EAR motif conservation in GL2 homologs across monocots and dicots.

**Figure S4.** The C-terminal EAR-motif mutants are defective in homodimerization in Y2H assays.

**Figure S5.** Western blotting confirms a comparable expression level of corepressors in Y2H assays.

**Figure S6.** Principal component analysis (PCA) of RNA-seq data.

**Figure S7.** Tissue-specific expression profiles of the *GL2* and *TPL/TPR* genes.

**Table S1.** Putative EAR motifs in HD-Zip IV and HD-Zip III proteins from Arabidopsis.

**Table S2.** TPL and GL2 EAR motif interaction report.

**Table S3.** Oligonucleotide primers used in this study.

**Dataset S1 (Separate sheet is xls file).** List of DEGs that are down-regulated in *gl2* compared to wild type (padj<0.05, log2FC>1).

**Dataset S2 (Separate sheet is xls file).** List of DEGs that are up-regulated in *gl2* compared to wild type (padj<0.05, log2FC>1).

**Dataset S3 (Separate sheet is xls file).** List of DEGs that are down-regulated in *gl2^ΔEAR-N^* compared to wild type (padj<0.05, log2FC>1).

**Dataset S4 (Separate sheet is xls file).** List of DEGs that are up-regulated in *gl2^ΔEAR-N^* compared to wild type (padj<0.05, log2FC>1).

**Dataset S5 (Separate sheet is xls file).** List of DEGs that are up-regulated in *gl2^ΔEAR-N^* compared to *gl2* (padj<0.05, log2FC>1).

**Dataset S6 (Separate sheet is xls file).** List of DEGs that are down-regulated in *gl2^ΔEAR-N^* compared to *gl2* (padj<0.05, log2FC>1).

## References

1. Ahmad, B., Lerma-Reyes, R., Mukherjee, T., Nguyen, H.V., Weber, A.L., Cummings, E.E., Schulze, W.X., Comer, J.R., and Schrick, K. (2024). Nuclear localization of Arabidopsis HD-Zip IV transcription factor GLABRA2 is driven by Importin α. *Journal of Experimental Botany*, erae326.

2. Baile, F., Merini, W., Hidalgo, I., and Calonje, M. (2021). EAR domain-containing transcription factors trigger PRC2-mediated chromatin marking in Arabidopsis. The Plant Cell 33, 2701–2715.

3. Bieluszewski, T., Xiao, J., Yang, Y., and Wagner, D. (2021). PRC2 activity, recruitment, and silencing: a comparative perspective. Trends in Plant Science 26, 1186–1198.

4. Cai, Y., Zhang, Y., Loh, Y.P., Tng, J.Q., Lim, M.C., Cao, Z., Raju, A., Lieberman Aiden, E., Li, S., and Manikandan, L. (2021). H3K27me3-rich genomic regions can function as silencers to repress gene expression via chromatin interactions. Nature Communications 12, 719.

5. Chanvivattana, Y., Bishopp, A., Schubert, D., Stock, C., Moon, Y.-H., Sung, Z.R., and Goodrich, J. (2004). Interaction of Polycomb-group proteins controlling flowering in Arabidopsis. Development 131, 5263–5276.

6. Cheng, J., Wang, J., Bi, S., Li, M., Wang, L., Wang, L., Li, T., Zhang, X., Gao, Y., and Zhu, L. (2024). GLABRA 2 regulates ETHYLENE OVERPRODUCER 1 accumulation during nutrient deficiency-induced root hair growth. Plant Physiology 195, 1906–1924.

7. Chew, W., Hrmova, M., and Lopato, S. (2013). Role of homeodomain leucine zipper (HD-Zip) IV transcription factors in plant development and plant protection from deleterious environmental factors. International Journal of Molecular Sciences 14, 8122–8147.

8. Chow, V., Kirzinger, M.W., and Kagale, S. (2023). Lend me your ears: a systematic review of the broad functions of ear motif-containing transcriptional repressors in plants. Genes 14, 270.

9. Clough, S.J., and Bent, A.F. (1998). Floral dip: a simplified method for Agrobacterium-mediated transformation of *Arabidopsis thaliana*. The Plant Journal 16, 735–743.

10. De Vries, S.J., Van Dijk, M., and Bonvin, A.M. (2010). The HADDOCK web server for data-driven biomolecular docking. Nature Protocols 5, 883–897.

11. Deka, B., and Singh, K.K. (2017). Multifaceted regulation of gene expression by the apoptosis-and splicing-associated protein complex and its components. International Journal of Biological Sciences 13, 545.

12. Di Cristina, M., Sessa, G., Dolan, L., Linstead, P., Baima, S., Ruberti, I., and Morelli, G. (1996). The Arabidopsis Athb-10 (GLABRA2) is an HD-Zip protein required for regulation of root hair development. The Plant Journal 10, 393–402.

13. Eric F. Pettersen, T.D.G., Conrad C. Huang, Gregory S. Couch, Daniel M. Greenblatt, Elaine C. Meng, Thomas E. Ferrin. (2004). UCSF Chimera—A visualization system for exploratory research and analysis. Journal of Computational Chemistry 25.13, 1605–1612.

14. Gietz, R.D., and Woods, R.A. (2002). Transformation of yeast by lithium acetate/single-stranded carrier DNA/polyethylene glycol method. In Methods in Enzymology (Elsevier), pp. 87–96.

15. Guan, X.Y., Li, Q.J., Shan, C.M., Wang, S., Mao, Y.B., Wang, L.J., and Chen, X.Y. (2008). The HD-Zip IV gene GaHOX1 from cotton is a functional homologue of the Arabidopsis GLABRA2. Physiologia Plantarum 134, 174–182.

16. He, J.-X., Gendron, J.M., Sun, Y., Gampala, S.S., Gendron, N., Sun, C.Q., and Wang, Z.-Y. (2005). BZR1 is a transcriptional repressor with dual roles in brassinosteroid homeostasis and growth responses. Science 307, 1634–1638.

17. Hill, K., Wang, H., and Perry, S.E. (2008). A transcriptional repression motif in the MADS factor AGL15 is involved in recruitment of histone deacetylase complex components. The Plant Journal 53, 172–185.

18. Hiratsu, K., Matsui, K., Koyama, T., and Ohme-Takagi, M. (2003). Dominant repression of target genes by chimeric repressors that include the EAR motif, a repression domain, in Arabidopsis. The Plant Journal 34, 733–739.

19. Ito, M., Sentoku, N., Nishimura, A., Hong, S.K., Sato, Y., and Matsuoka, M. (2002). Position dependent expression of GL2-type homeobox gene, Roc1: significance for protoderm differentiation and radial pattern formation in early rice embryogenesis. The Plant Journal 29, 497–507.

20. Ito, M., Sentoku, N., Nishimura, A., Hong, S.-K., Sato, Y., and Matsuoka, M. (2003). Roles of Rice GL2-type Homeobox Genes in Epidermis Differentiation. Breeding Sci 53, 245–253.

21. Johnston, A.J., Kirioukhova, O., Barrell, P.J., Rutten, T., Moore, J.M., Baskar, R., Grossniklaus, U., and Gruissem, W. (2010). Dosage-sensitive function of retinoblastoma related and convergent epigenetic control are required during the Arabidopsis life cycle. PLoS Genetics 6, e1000988.

22. Kagale, S., and Rozwadowski, K. (2011). EAR motif-mediated transcriptional repression in plants: an underlying mechanism for epigenetic regulation of gene expression. Epigenetics 6, 141–146.

23. Kagale, S., Links, M.G., and Rozwadowski, K. (2010). Genome-wide analysis of ethylene-responsive element binding factor-associated amphiphilic repression motif-containing transcriptional regulators in Arabidopsis. Plant Physiology 152, 1109–1134.

24. Khosla, A., Paper, J.M., Boehler, A.P., Bradley, A.M., Neumann, T.R., and Schrick, K. (2014). HD-Zip proteins GL2 and HDG11 have redundant functions in Arabidopsis trichomes, and GL2 activates a positive feedback loop via MYB23. The Plant Cell 26, 2184–2200.

25. Lamiable, A., Thévenet, P., Rey, J., Vavrusa, M., Derreumaux, P., and Tufféry, P. (2016). PEP-FOLD3: faster de novo structure prediction for linear peptides in solution and in complex. Nucleic Acids Research 44, W449–W454.

26. Lee, M.-S., An, J.-H., and Cho, H.-T. (2016). Biological and molecular functions of two EAR motifs of Arabidopsis IAA7. Journal of Plant Biology 59, 24–32.

27. Lin, Q., Ohashi, Y., Kato, M., Tsuge, T., Gu, H., Qu, L.-J., and Aoyama, T. (2015). GLABRA2 directly suppresses basic helix-loop-helix transcription factor genes with diverse functions in root hair development. The Plant Cell 27, 2894–2906.

28. Liu, X., Yang, S., Zhao, M., Luo, M., Yu, C.-W., Chen, C.-Y., Tai, R., and Wu, K. (2014). Transcriptional repression by histone deacetylases in plants. Molecular Plant 7, 764–772.

29. Long, J.A., Ohno, C., Smith, Z.R., and Meyerowitz, E.M. (2006). TOPLESS regulates apical embryonic fate in Arabidopsis. Science 312, 1520–1523.

30. Long, J.A., Woody, S., Poethig, S., Meyerowitz, E.M., and Barton, M.K. (2002). Transformation of shoots into roots in Arabidopsis embryos mutant at the TOPLESS locus. Development 129, 2797–2806.

31. Martin-Arevalillo, R., Nanao, M.H., Larrieu, A., Vinos-Poyo, T., Mast, D., Galvan-Ampudia, C., Brunoud, G., Vernoux, T., Dumas, R., and Parcy, F. (2017). Structure of the Arabidopsis TOPLESS corepressor provides insight into the evolution of transcriptional repression. Proceedings of the National Academy of Sciences U.S.A. 114, 8107–8112.

32. Masucci, J.D., Rerie, W.G., Foreman, D.R., Zhang, M., Galway, M.E., Marks, M.D., and Schiefelbein, J.W. (1996). The homeobox gene GLABRA 2 is required for position-dependent cell differentiation in the root epidermis of *Arabidopsis thaliana*. Development 122, 1253–1260.

33. Mozgova, I., Kohler, C., and Hennig, L. (2015). Keeping the gate closed: functions of the polycomb repressive complex PRC2 in development. Plant Journal 83, 121–132.

34. Navarro, M.A., Navarro, C., Hernández, L.E., Garnica, M., Franco-Zorrilla, J.M., Burko, Y., González-Serrano, S., García-Mina, J.M., Pruneda-Paz, J., and Chory, J. (2024). GLABRA 2 transcription factor integrates arsenic tolerance with epidermal cell fate determination. New Phytologist 244, 1882–1900.

35. Oh, E., Zhu, J.-Y., Ryu, H., Hwang, I., and Wang, Z.-Y. (2014). TOPLESS mediates brassinosteroid-induced transcriptional repression through interaction with BZR1. Nature Communications 5, 4140.

36. Ohashi, Y., Oka, A., Rodrigues-Pousada, R., Possenti, M., Ruberti, I., Morelli, G., and Aoyama, T. (2003). Modulation of phospholipid signaling by GLABRA2 in root-hair pattern formation. Science 300, 1427–1430.

37. Ohta, M., Matsui, K., Hiratsu, K., Shinshi, H., and Ohme-Takagi, M. (2001). Repression domains of class II ERF transcriptional repressors share an essential motif for active repression. The Plant Cell 13, 1959–1968.

38. Plant, A.R., Larrieu, A., and Causier, B. (2021). Repressor for hire! The vital roles of TOPLESS-mediated transcriptional repression in plants. New Phytologist 231, 963–973.

39. Qüesta, J.I., Song, J., Geraldo, N., An, H., and Dean, C. (2016). Arabidopsis transcriptional repressor VAL1 triggers Polycomb silencing at FLC during vernalization. Science 353, 485–488.

40. Rerie, W.G., Feldmann, K.A., and Marks, M.D. (1994). The GLABRA2 gene encodes a homeo domain protein required for normal trichome development in Arabidopsis. Genes & Development 8, 1388–1399.

41. Shan, C.-M., Shangguan, X.-X., Zhao, B., Zhang, X.-F., Chao, L.-m., Yang, C.-Q., Wang, L.-J., Zhu, H.-Y., Zeng, Y.-D., and Guo, W.-Z. (2014). Control of cotton fibre elongation by a homeodomain transcription factor GhHOX3. Nature Communications 5, 5519.

42. Shen, B., Sinkevicius, K.W., Selinger, D.A., and Tarczynski, M.C. (2006). The homeobox gene GLABRA2 affects seed oil content in Arabidopsis. Plant Molecular Biology 60, 377–387.

43. Shi, L., Katavic, V., Yu, Y., Kunst, L., and Haughn, G. (2012). Arabidopsis glabra2 mutant seeds deficient in mucilage biosynthesis produce more oil. The Plant Journal 69, 37–46.

44. Shu, J., Chen, C., Thapa, R.K., Bian, S., Nguyen, V., Yu, K., Yuan, Z.C., Liu, J., Kohalmi, S.E., and Li, C. (2019). Genome-wide occupancy of histone H3K27 methyltransferases CURLY LEAF and SWINGER in Arabidopsis seedlings. Plant Direct 3, e00100.

45. Song, C.-P., and Galbraith, D.W. (2006). AtSAP18, an orthologue of human SAP18, is involved in the regulation of salt stress and mediates transcriptional repression in Arabidopsis. Plant Molecular Biology 60, 241–257.

46. Sun, Y., Fan, X.-Y., Cao, D.-M., Tang, W., He, K., Zhu, J.-Y., He, J.-X., Bai, M.-Y., Zhu, S., and Oh, E. (2010). Integration of brassinosteroid signal transduction with the transcription network for plant growth regulation in Arabidopsis. Developmental Cell 19, 765–777.

47. Szemenyei, H., Hannon, M., and Long, J.A. (2008). TOPLESS mediates auxin-dependent transcriptional repression during Arabidopsis embryogenesis. Science 319, 1384–1386.

48. Tao, Q., Guo, D., Wei, B., Zhang, F., Pang, C., Jiang, H., Zhang, J., Wei, T., Gu, H., and Qu, L.-J. (2013). The TIE1 transcriptional repressor links TCP transcription factors with TOPLESS/TOPLESS-RELATED corepressors and modulates leaf development in Arabidopsis. The Plant Cell 25, 421–437.

49. Tao, Z., Zhu, L., Li, H., Sun, B., Liu, X., Li, D., Hu, W., Wang, S., Miao, X., and Shi, Z. (2024). ACL1-ROC4/5 complex reveals a common mechanism in rice response to brown planthopper infestation and drought. Nature Communications 15, 8107.

50. Vernoud, V., Laigle, G., Rozier, F., Meeley, R.B., Perez, P., and Rogowsky, P.M. (2009). The HD-ZIP IV transcription factor OCL4 is necessary for trichome patterning and anther development in maize. The Plant Journal 59, 883–894.

51. Wang, J., Alekseenko, A., Kozakov, D., and Miao, Y. (2019). Improved modeling of peptide-protein binding through global docking and accelerated molecular dynamics simulations. Frontiers in Molecular Biosciences 6, 112.

52. Wang, L., Kim, J., and Somers, D.E. (2013). Transcriptional corepressor TOPLESS complexes with pseudoresponse regulator proteins and histone deacetylases to regulate circadian transcription. Proceedings of the National Academy of Sciences U.S.A 110, 761–766.

53. Wang, W., Wang, X., Wang, Y., Zhou, G., Wang, C., Hussain, S., Adnan, Lin, R., Wang, T., and Wang, S. (2020a). SlEAD1, an EAR motif-containing ABA down-regulated novel transcription repressor regulates ABA response in tomato. GM Crops & Food 11, 275–289.

54. Wang, X., Wang, X., Hu, Q., Dai, X., Tian, H., Zheng, K., Wang, X., Mao, T., Chen, J.G., and Wang, S. (2015). Characterization of an activation-tagged mutant uncovers a role of GLABRA 2 in anthocyanin biosynthesis in *Arabidopsis*. The Plant Journal 83, 300–311.

55. Wang, X., Bi, S., Wang, L., Li, H., Gao, B.-a., Huang, S., Qu, X., Cheng, J., Wang, S., and Liu, C. (2020b). GLABRA2 regulates actin bundling protein VILLIN1 in root hair growth in response to osmotic stress. Plant Physiology 184, 176–193.

56. Wang, Z., Tian, X., Zhao, Q., Liu, Z., Li, X., Ren, Y., Tang, J., Fang, J., Xu, Q., and Bu, Q. (2018). The E3 ligase DROUGHT HYPERSENSITIVE negatively regulates cuticular wax biosynthesis by promoting the degradation of transcription factor ROC4 in rice. The Plant Cell 30, 228–244.

57. Western, T.L., Burn, J., Tan, W.L., Skinner, D.J., Martin-McCaffrey, L., Moffatt, B.A., and Haughn, G.W. (2001). Isolation and characterization of mutants defective in seed coat mucilage secretory cell development in Arabidopsis. Plant Physiology 127, 998–1011.

58. Wu, M., Chang, J., Han, X., Shen, J., Yang, L., Hu, S., Huang, B.-B., Xu, H., Xu, M., and Wu, S. (2023). A HD-ZIP transcription factor specifies fates of multicellular trichomes via dosage-dependent mechanisms in tomato. Developmental Cell 58, 278–288. e275.

59. Wu, R., and Citovsky, V. (2017a). Adaptor proteins GIR1 and GIR2. I. Interaction with the repressor GLABRA2 and regulation of root hair development. Biochemical Biophysical Research Communications 488, 547–553.

60. Wu, R., and Citovsky, V. (2017b). Adaptor proteins GIR1 and GIR2. II. Interaction with the co-repressor TOPLESS and promotion of histone deacetylation of target chromatin. Biochem Biophys Res Commun 488, 609–613.

61. Xiao, J., Jin, R., Yu, X., Shen, M., Wagner, J.D., Pai, A., Song, C., Zhuang, M., Klasfeld, S., and He, C. (2017). Cis and trans determinants of epigenetic silencing by Polycomb repressive complex 2 in Arabidopsis. Nature Genetics 49, 1546–1552.

62. Xu, Y., Kong, W., Wang, F., Wang, J., Tao, Y., Li, W., Chen, Z., Fan, F., Jiang, Y., and Zhu, Q.H. (2021). Heterodimer formed by ROC8 and ROC5 modulates leaf rolling in rice. Plant Biotechnology Journal 19, 2662–2672.

63. Yang, J., Liu, Y., Yan, H., Tian, T., You, Q., Zhang, L., Xu, W., and Su, Z. (2018). PlantEAR: functional analysis platform for plant EAR motif-containing proteins. Frontiers in Genetics 9, 590.

64. Zhang, Y., Iratni, R., Erdjument-Bromage, H., Tempst, P., and Reinberg, D. (1997). Histone deacetylases and SAP18, a novel polypeptide, are components of a human Sin3 complex. Cell 89, 357–364.

65. Zhu, Z., Xu, F., Zhang, Y., Cheng, Y.T., Wiermer, M., Li, X., and Zhang, Y. (2010). Arabidopsis resistance protein SNC1 activates immune responses through association with a transcriptional corepressor. Proceedings of the National Academy of Sciences 107, 13960–13965.

66. Zou, L.-p., Sun, X.-h., Zhang, Z.-g., Liu, P., Wu, J.-x., Tian, C.-j., Qiu, J.-l., and Lu, T.-g. (2011). Leaf rolling controlled by the homeodomain leucine zipper class IV gene Roc5 in rice. Plant Physiology 156, 1589–1602.

