## Supplemental Figures and Tables for "GLABRA2 regulates gene expression via its own EAR-motif mediated recruitment of the TPL/TPR corepressors"

**List of Supplemental Figures, Tables, and Datasets:**

**Figure S1.** Suppression of *gl2* trichome defects by *gir1* mutation.

**Figure S2.** GL2 fails to interact with GIR1 and GIR2 in yeast two-hybrid (Y2H) assays.

**Figure S3.** EAR motif conservation in GL2 homologs across monocots and dicots.

**Figure S4.** The C-terminal EAR-motif mutants are defective in homodimerization in Y2H assays.

**Figure S5.** Western blotting confirms a comparable expression level of corepressors in Y2H assays.

**Figure S6.** Principal component analysis (PCA) of RNA-seq data.

**Figure S7.** Tissue-specific expression profiles of the *GL2* and *TPL/TPR* genes.

**Table S1.** Putative EAR motifs in HD-Zip IV and HD-Zip III proteins from Arabidopsis.

**Table S2.** TPL and GL2 EAR motif interaction report.

**Table S3.** Oligonucleotide primers used in this study.

**Dataset S1 (Separate sheet is xls file).** List of DEGs that are down-regulated in *gl2* compared to wild type (padj<0.05, log2FC>1).

**Dataset S2 (Separate sheet is xls file).** List of DEGs that are up-regulated in *gl2* compared to wild type (padj<0.05, log2FC>1).

**Dataset S3 (Separate sheet is xls file).** List of DEGs that are down-regulated in *gl2*$^{\Delta EAR\text{-}N}$ compared to wild type (padj<0.05, log2FC>1).

**Dataset S4 (Separate sheet is xls file).** List of DEGs that are up-regulated in *gl2*$^{\Delta EAR\text{-}N}$ compared to wild type (padj<0.05, log2FC>1).

**Dataset S5 (Separate sheet is xls file).** List of DEGs that are up-regulated in *gl2*$^{\Delta EAR\text{-}N}$ compared to *gl2* (padj<0.05, log2FC>1).

**Dataset S6 (Separate sheet is xls file).** List of DEGs that are down-regulated in *gl2*$^{\Delta EAR\text{-}N}$ compared to *gl2* (padj<0.05, log2FC>1).

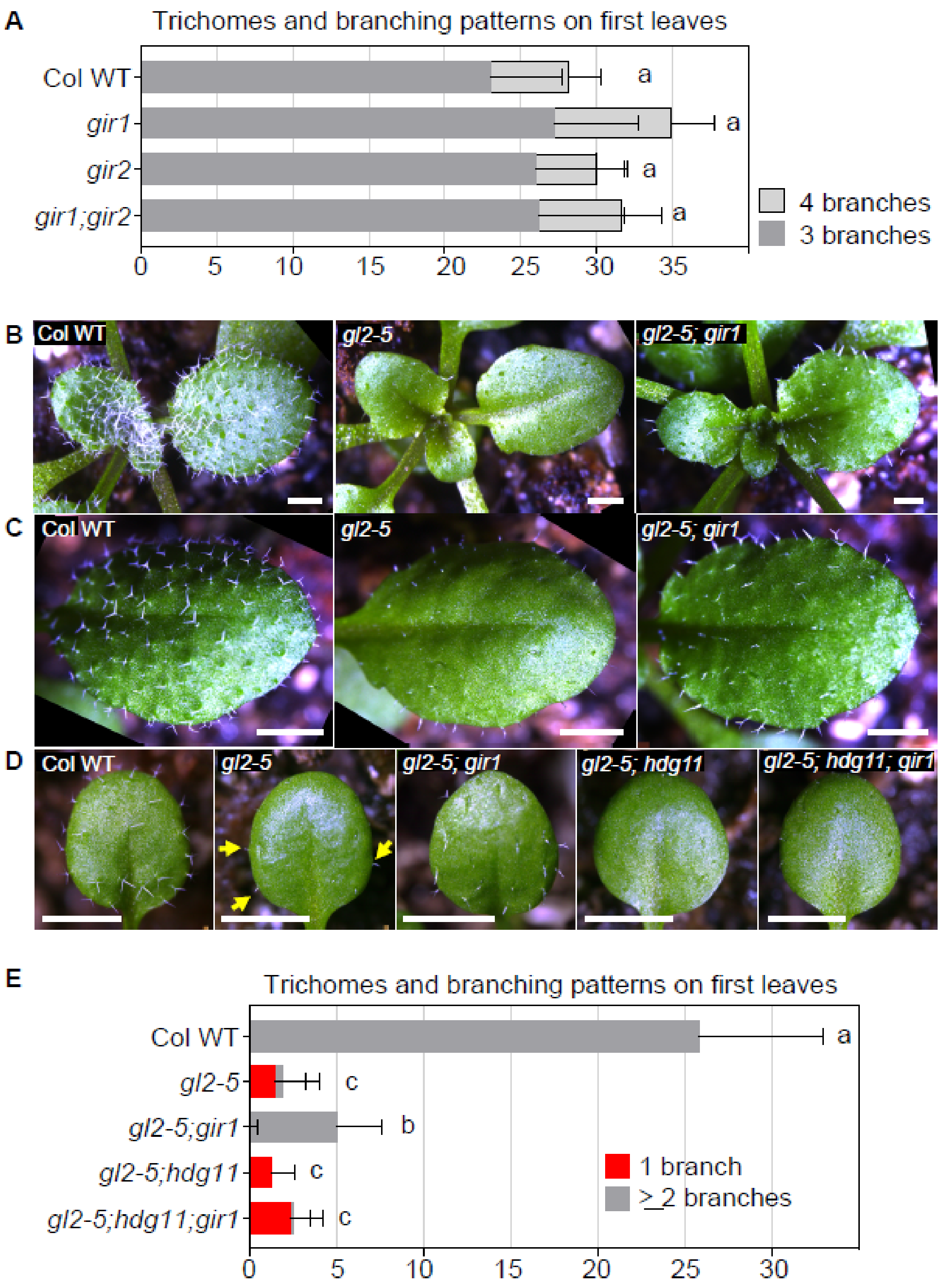

**Figure S1. Suppression of *gl2* trichome defects by *gir1* mutation.**

**(A)** Quantification of trichome numbers and branching on first leaves. The *gir1*, *gir2* and *gir1;gir2* double mutants display trichome phenotypes indistinguishable from Columbia wild type (Col WT). Error bars depict the SD, and letters denote no significant differences between genotypes for n=20 plants as determined by one-way ANOVA and Tukey's test ($p < 0.05$).

**(B and C)** Leaf trichome phenotypes indicate that the glabrous phenotype of *gl2-5* mutants is partially suppressed by *gir1* in *gl2-5;gir1* double mutants. Bar = 2 mm.
**(D)** Trichome phenotypes on first leaves. The trichome defects of *gl2-5* mutants appear suppressed in the *gl2-5;gir1* double mutant. In the *gl2-5;gir1;hdg11* triple mutant, the trichome defects appear similar to that of *gl2-5* and *gl2-5;hdg11*. Bar = 1 mm
**(E)** Quantification of trichome numbers and branching on first leaves for the genotypes shown in **(D)**. Error bars depict the SD, and letters denote significant differences between genotypes for n=20 plants as determined by one-way ANOVA and Tukey's test ($p < 10^{-5}$).

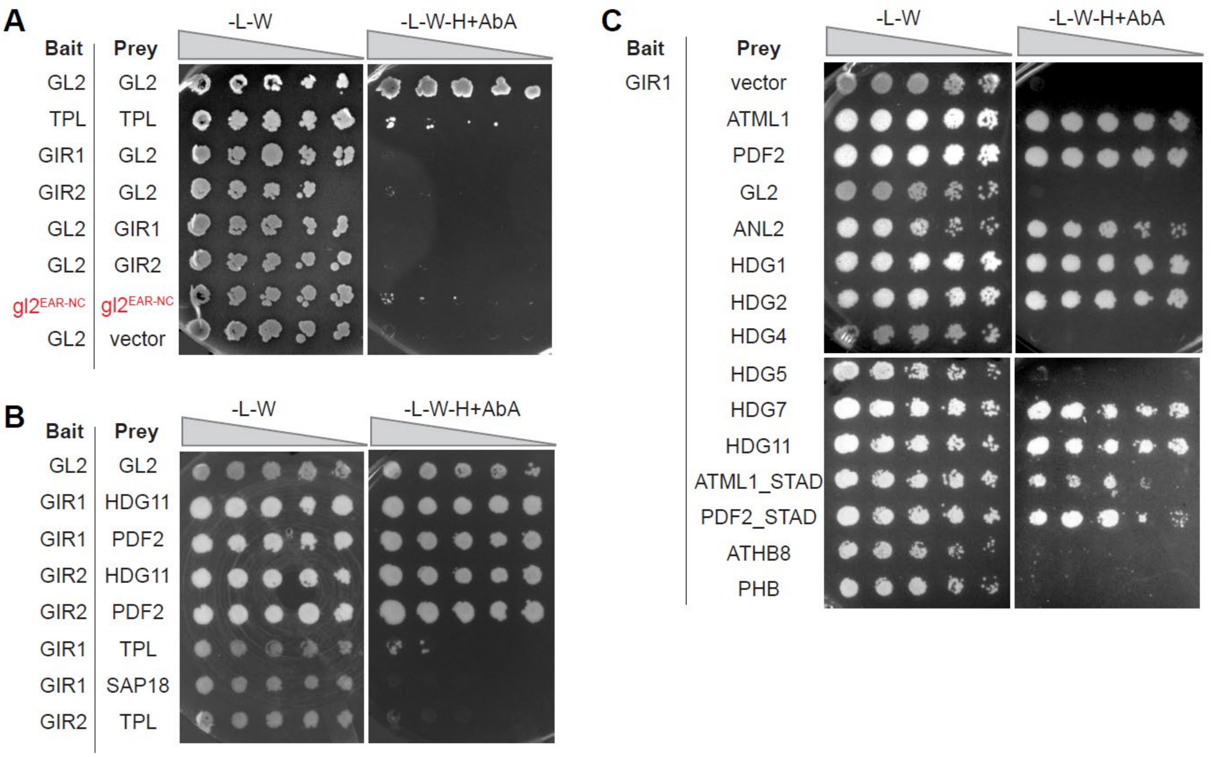

**Figure S2. GL2 fails to interact with GIR1 and GIR2 in yeast two-hybrid (Y2H) assays.**

**(A-C)** Y2H Bait and prey proteins are indicated on the left. Growth on -L-W signifies presence of both plasmids, and growth on selective media (-L-W-H+AbA) indicates protein interaction. Four-fold dilutions of yeast are indicated left to right.

**(A)** Y2H assays indicate that GL2 interacts with itself but not with GIR1 or GIR2. GL1 lack of interaction with the empty vector serves as a negative control.

**(B)** GIR1 and GIR2 interact with the HD-Zip IV transcription factors HDG11, PDF2, and ATML1. The GL2-GL2 self-interaction served as a positive control.

**(C)** GIR1 shows specificity of interaction with a subset of HD-Zip IV transcription factors. including ATML1, PDF2, ANL2, HDG1, HDG2, HDG7, and HDG11. Additionally, GIR1 was observed to interact with the START Adjacent Domain (STAD) from ATML1 and PDF2. GIR1 failed to interact with GL2, HDG4, HDG5, and the HD-Zip IV transcription factors ATHB8 and PHB.

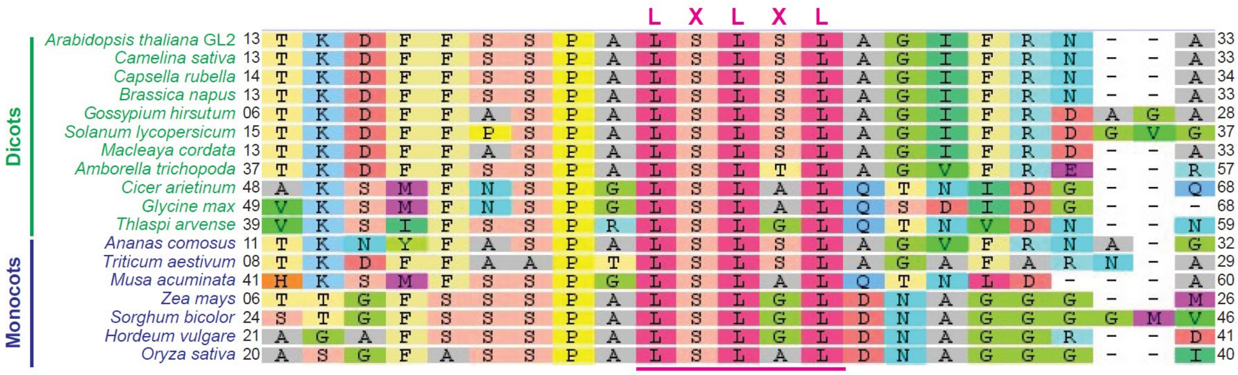

**Figure S3. EAR motif conservation in GL2 homologs across monocots and dicots.**

Multiple protein sequence alignment demonstrates the conservation of the N-terminal EAR motif (LxLxL) in *Arabidopsis thaliana* GL2 (At1g79840) and its homologs in monocotyledonous and dicotyledonous plants. The alignment was performed using Geneious 7.1.9 (https://www.geneious.com) with the Geneious alignment algorithm and default settings (global alignment with free end gaps and cost matrix BLOSUM62). The sequence and the corresponding initial and terminal residue numbers are provided. Accession IDs: *Arabidopsis thaliana* GL2 (NP_565223.1), *Camelina sativa* (XP_010430051.1), Capsella rubella (XP_023642973.1), *Brassica napus* (XP_013680822), *Gossypium hirsutum* (NP_001314068.1), *Solanum lycopersicum* (XP_004235676.1), *Macleaya cordata* (OVA20332.1), *Amborella trichopoda* (XP_020518333.1), Cicer arietinum (XP_004510857.1), *Glycine max* (KAG5014136.1), *Thlaspi arvense* (CAH2057809.1), *Musa acuminata* (XP_009401771.1), *Ananas comosus* (XP_020098513.1), *Triticum aestivum* (XP_044341402.1), *Zea mays* (NP_001105493.1), *Sorghum bicolor* (XP_002457484.1), *Hordeum vulgare* (XP_044950934.1), and *Oryza sativa* (NP_001403749.1).

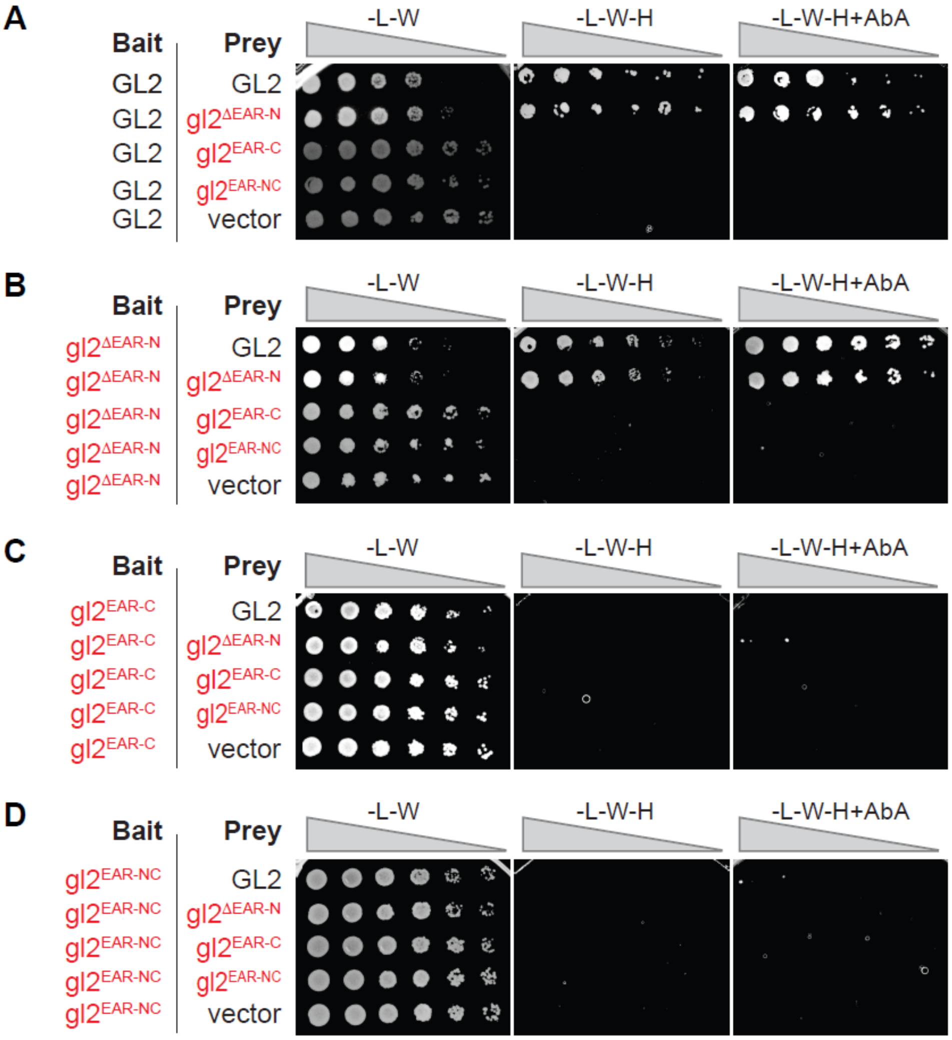

**Figure S4. The C-terminal EAR-motif mutants are defective in homodimerization in Y2H assays.**

**(A)** Y2H homodimerization assays reveal that GL2 dimerizes with itself and $gl2^{\Delta EAR\text{-}N}$, but not with $gl2^{EAR\text{-}C}$ or $gl2^{EAR\text{-}NC}$.

**(B)** $gl2^{\Delta EAR\text{-}N}$ dimerizes with itself and wild-type GL2, but not with $gl2^{EAR\text{-}C}$ or $gl2^{EAR\text{-}NC}$.

**(C and D)** $gl2^{EAR\text{-}C}$ **(C)** and $gl2^{EAR\text{-}NC}$ **(D)** fail to dimerize with wild-type GL2 or the EAR-motif mutants. The permissive growth condition is shown in the left panel (-L-W), while homodimerization is indicated in the panels depicting selective media (-L-W-H and -L-W-H+AbA). The empty vector served as a negative control. Increasing four-fold dilutions are indicated from left to right.

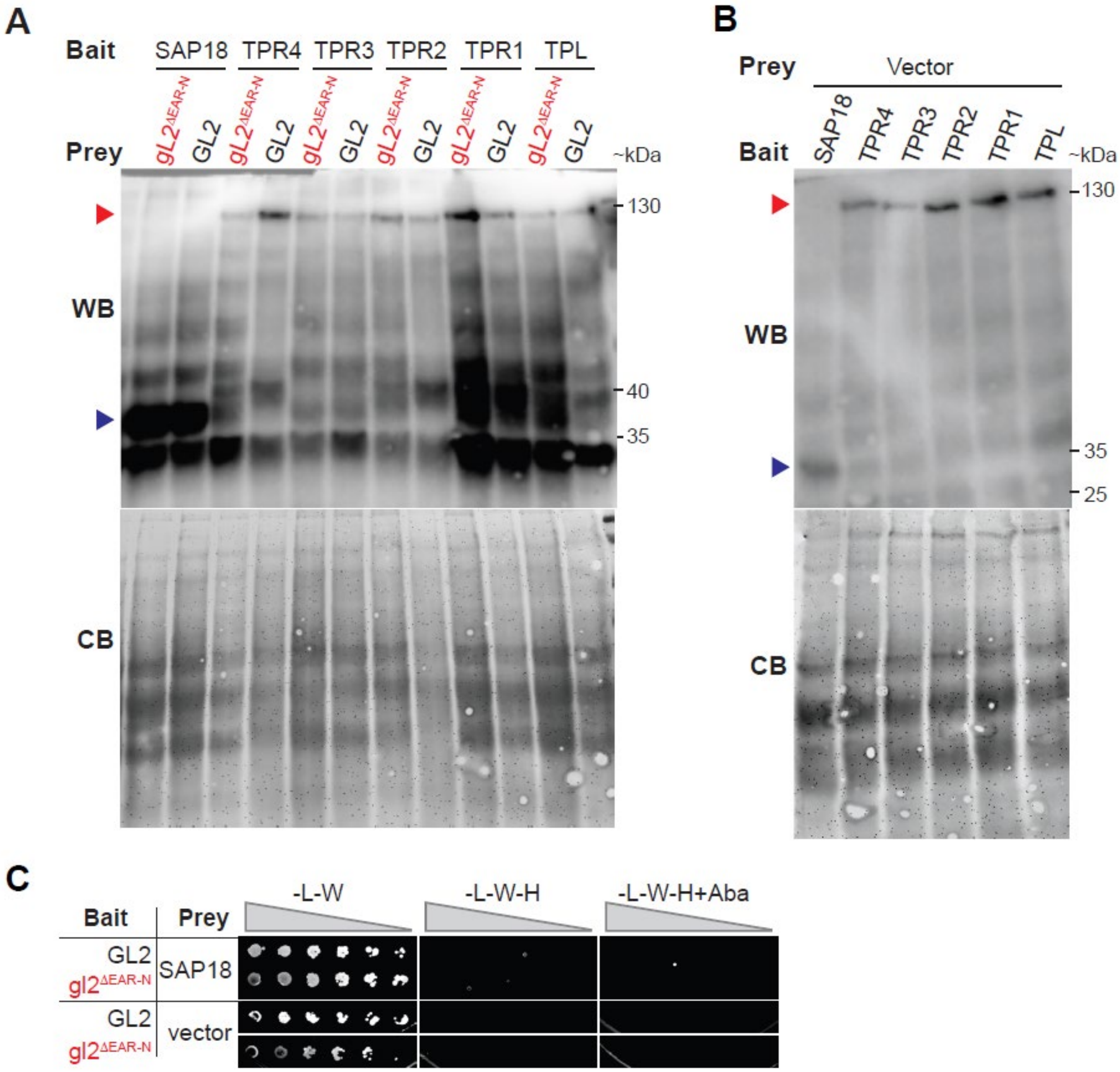

**Figure S5. Western blotting confirms a comparable expression level of corepressors in Y2H assays.** Related to Fig. 3E.

**(A and B)** Bait proteins fused to GAL4 (DBD) were resolved by SDS-PAGE followed by detection with anti-GAL4 Ab. Red arrowhead indicates expression of TPL and TPR proteins when used as baits with GL2 or gl2$^{\Delta EAR\text{-}N}$ **(A)** or vector **(B)** as prey; blue arrowhead indicates expression of SAP18. The Coomassie blue (CB) stained blots are shown at the bottom to indicate total protein loading. Approximate kDa protein sizes are given on the right.

**(C)** Y2H assays show that SAP18 fails to interact with GL2 when used as prey. GL2 and gl2$^{\Delta EAR\text{-}N}$ fused with GAL4 (DBD) served as baits, and SAP18 fused with GAL4 (AD) served as prey. Interactions are indicated on selective media lacking Leu, Trp, and His (-L-W-H), in the presence (right) or absence (middle) of Aureobasidin A (AbA). GL2 failed to show interaction with the empty vector negative control. Increasing four-fold dilutions are indicated from left to right.

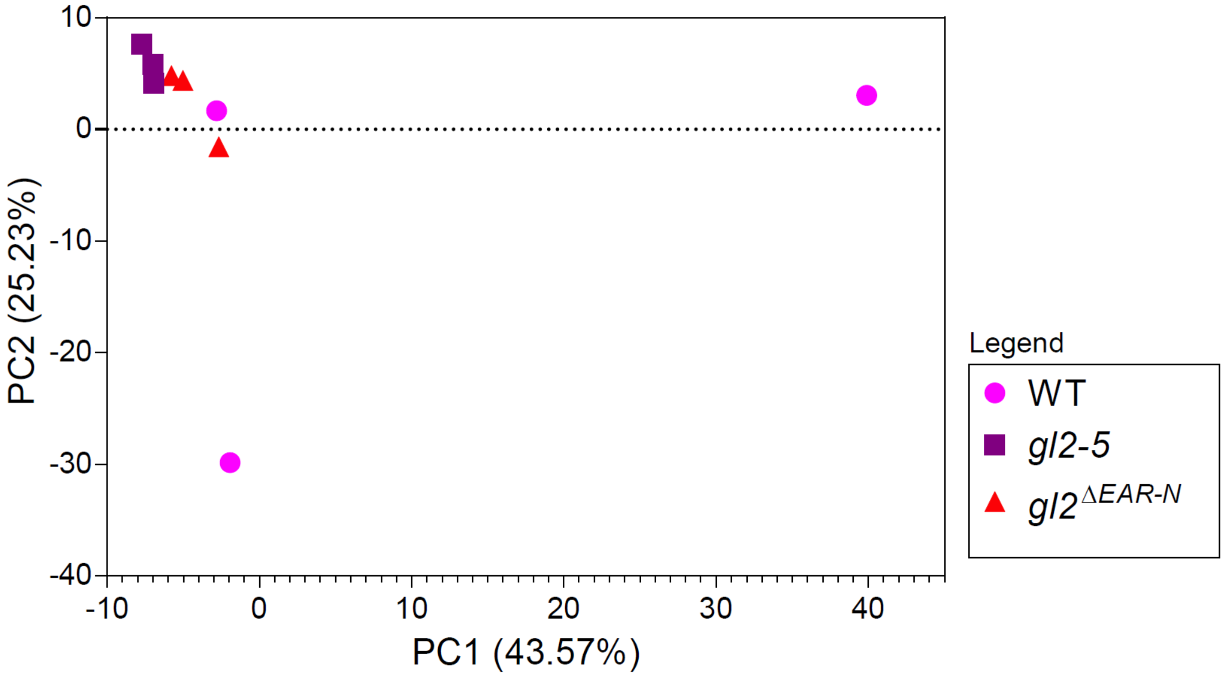

**Figure S6. Principal component analysis (PCA) of RNA-seq data.**

Three biological replicates of each genotype were subject to principal component analysis (PCA) performed on $log_2$(FPKM+1) expression values across all genes for each sample. Symbols (circles, squares, triangles) represent biological replicates, indicated by genotype (wild type (WT), *gl2*, *gl2*$^{\Delta EAR\text{-}N}$). Axes show PC1 and PC2 with the percentage of variance explained (PC1 = 43.57%, PC2 = 25.23% in this dataset). Greater distances indicate lower transcriptome similarity. WT samples separate strongly along PC1, while *gl2*$^{\Delta EAR\text{-}N}$ is positioned between WT and *gl2-5*, consistent with an intermediate transcriptional state.

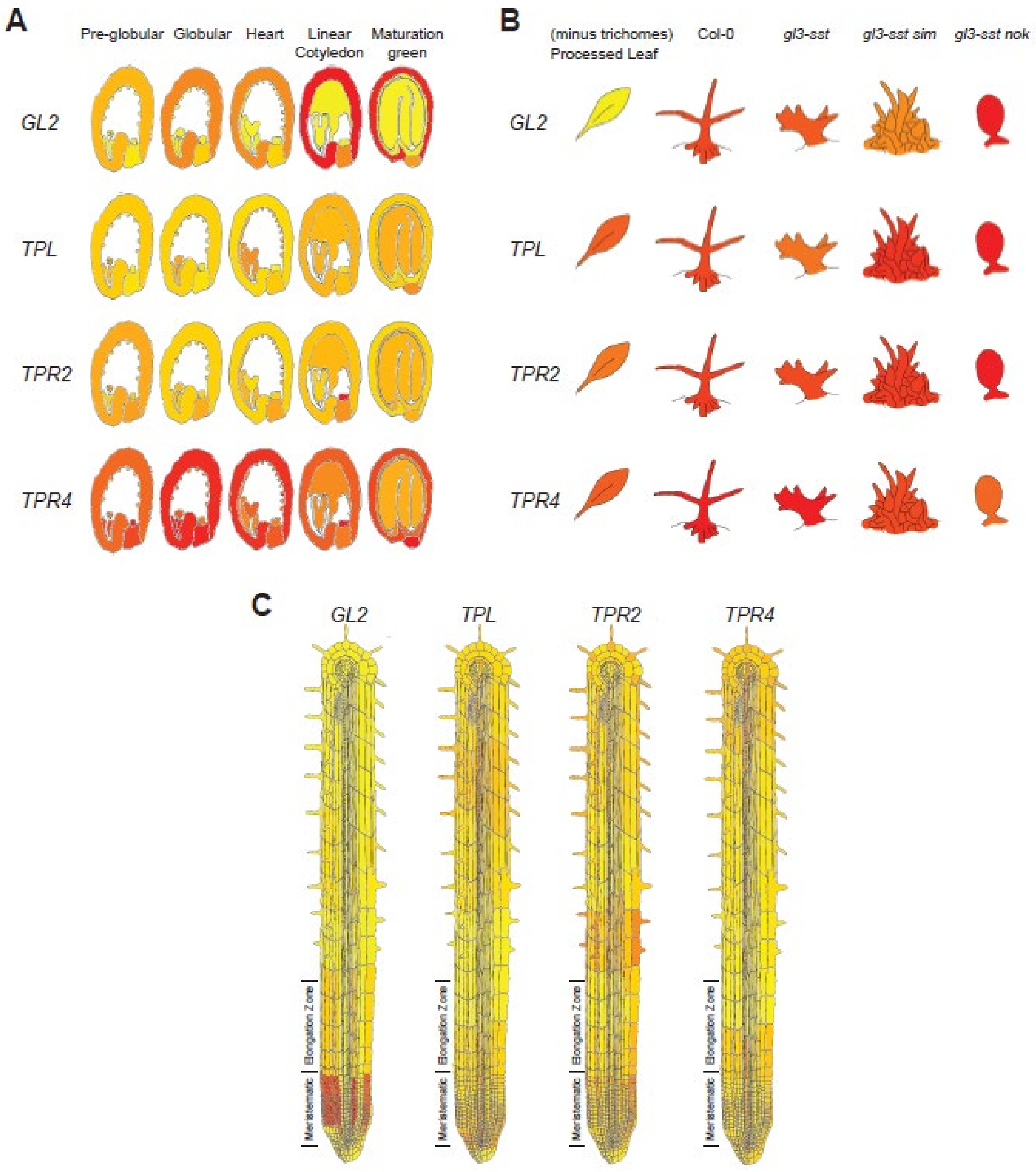

**Figure S7. Tissue-specific expression profiles of the *GL2* and *TPL/TPR* genes.**
Publicly available mRNA expression data for GL2, TPL, TPR2, and TPR4 were retrieved from the Arabidopsis eFP Browser (https://bar.utoronto.ca/efp//cgi-bin/efpWeb.cgi). Color scale: red (maximum) to yellow (minimum).
**(A)** mRNA expression of GL2, TPL, TPR2, and TPR4 at various seed developmental stages. Data derived from (Le et al., 2010).

**(B)** Expression in leaves (minus trichomes) and trichomes from wild-type Columbia (Col) and mutant lines (*gl3-sst*, *gl3-sst sim*, and *gl3-sst nok*). Data derived from (Marks et al., 2009; Gilding and Marks, 2010).
**(C)** Root cell-type specific expression patterns in 7-day-old seedlings. Data derived from (Brady et al., 2007).

**Table S1**. **Putative EAR motifs in HD-Zip IV and HD-Zip III proteins from Arabidopsis.** Sequences and positions of putative EAR motifs are mapped to protein domains. GL2 contains two putative EAR motifs, one at the N-terminus and the other within the STAD C-terminal domain. A putative EAR motif is also found in the charophycean green algae *Spirogyra pratensis* HD-Zip IV (GenBank: WOR75494.1). For comparison, no putative EAR motifs were detected in the Arabidopsis HD-Zip III proteins nor in the *Spirogyra pratensis* HD-Zip III (GenBank: ANH56717.1).

| **HD-Zip IV** | **Arabidopsis ID** | **N-terminus** | **HD** | **ZLZ** | **START** | **STAD** |
|---|---|---|---|---|---|---|
| GL2 | AT1G79840 | $_{22}$LSLSL$_{26}$ | - | - | - | $_{703}$LTLAL$_{707}$ |
| HDG1 | AT3G61150 | $_{53}$LSLGL$_{57}$ | - | - | - | - |
| ANL2 | AT4G00730 | $_{65}$LSLAL$_{69}$ | - | - | - | - |
| PDF2 | AT4G04890 | - | $_{98}$DLNLEP$_{103}$ | - | - | - |
| HDG3 | AT2G32370 | - | - | - | $_{281}$LALNL$_{285}$ | - |
| HDG5 | AT5G46880 | - | - | - | $_{436}$LLLVL$_{440}$ | - |
| HDG9 | AT5G17320 | - | - | - | - | $_{547}$LSLPL$_{551}$ |
| HDG10 | AT1G34650 | - | - | - | - | $_{538}$LSLPL$_{542}$ |
| ATML1 | AT4G21750 | - | - | - | - | - |
| HDG2 | AT1G05230 | - | - | - | - | - |
| HDG4 | AT4G17710 | - | - | - | - | - |
| FWA | AT4G25530 | - | - | - | - | - |
| HDG7 | AT5G52170 | - | - | - | - | - |
| HDG8 | AT3G03260 | - | - | - | - | - |
| HDG11 | AT1G73360 | - | - | | | |
| HDG12 | AT1G17920 | - | - | - | - | - |
| SpCH4 | *Spirogyra pratensis* | $_{49}$LDLAL$_{53}$ | - | | | |
| **HD-Zip III** | **Arabidopsis ID** | **N-terminus** | **HD** | **ZLZ** | **START** | **STAD** |
| AtHB-8 | AT4g32880 | - | - | - | - | - |
| CNA | AT1g52150 | - | - | - | - | - |
| PHB | AT2g34710 | - | - | - | - | - |
| PHV | AT1g30490 | - | - | - | - | - |
| REV | AT5g60690 | - | - | - | - | - |
| SpCH3 | *Spirogyra pratensis* | - | - | - | - | - |

**Table S2. TPL and GL2 EAR motif interaction report.**

The physicochemical properties of the amino acid interactions between the TPL 184 N-terminal residues and the GL2 EAR motif, determined using Discovery Studio (Version 2020, BIOVIA, USA), are provided. Related to **Figure 3D**.

| Name | Dist. (Å) | Category | Types | From | From Chemistry | To | To Chemistry |
|---|---|---|---|---|---|---|---|
| A:LYS71:HZ1 - B:GLY10:O | 1.64377 | Hydrogen Bond | Conventional Hydrogen Bond | A:LYS71:HZ1 | H-Donor | B:GLY10:O | H-Acceptor |
| A:LYS71:HZ3 - B:LEU6:O | 1.72166 | Hydrogen Bond | Conventional Hydrogen Bond | A:LYS71:HZ3 | H-Donor | B:LEU6:O | H-Acceptor |
| A:LYS78:HZ1 - B:SER5:OG | 1.90868 | Hydrogen Bond | Conventional Hydrogen Bond | A:LYS78:HZ1 | H-Donor | B:SER5:OG | H-Acceptor |
| A:ASN108:HD21 - B:ALA9:O | 2.17069 | Hydrogen Bond | Conventional Hydrogen Bond | A:ASN108:HD21 | H-Donor | B:ALA9:O | H-Acceptor |
| A:ASN108:HD22 - B:LEU8:O | 3.06757 | Hydrogen Bond | Conventional Hydrogen Bond | A:ASN108:HD22 | H-Donor | B:LEU8:O | H-Acceptor |
| A:LYS132:HN - B:SER1:OG | 2.85047 | Hydrogen Bond | Conventional Hydrogen Bond | A:LYS132:HN | H-Donor | B:SER1:OG | H-Acceptor |
| B:SER5:CB - A:GLU75:OE2 | 3.46794 | Hydrogen Bond | Carbon Hydrogen Bond | B:SER5:CB | H-Donor | A:GLU75:OE2 | H-Acceptor |
| B:ALA9:CB - A:PHE74 | 3.86511 | Hydrophobic | Pi-Sigma | B:ALA9:CB | C-H | A:PHE74 | Pi-Orbitals |
| A:LYS71 - B:LEU6 | 4.69039 | Hydrophobic | Alkyl | A:LYS71 | Alkyl | B:LEU6 | Alkyl |
| A:LEU111 - B:LEU8 | 5.20806 | Hydrophobic | Alkyl | A:LEU111 | Alkyl | B:LEU8 | Alkyl |
| B:PRO2 - A:ILE142 | 4.71973 | Hydrophobic | Alkyl | B:PRO2 | Alkyl | A:ILE142 | Alkyl |
| B:ALA9 - A:LEU111 | 4.44657 | Hydrophobic | Alkyl | B:ALA9 | Alkyl | A:LEU111 | Alkyl |
| A:LYS71:HZ1 - B:GLY10:O | 1.64377 | Hydrogen Bond | Conventional Hydrogen Bond | A:LYS71:HZ1 | H-Donor | B:GLY10:O | H-Acceptor |
| A:LYS71:HZ3 - B:LEU6:O | 1.72166 | Hydrogen Bond | Conventional Hydrogen Bond | A:LYS71:HZ3 | H-Donor | B:LEU6:O | H-Acceptor |
| A:LYS78:HZ1 - B:SER5:OG | 1.90868 | Hydrogen Bond | Conventional Hydrogen Bond | A:LYS78:HZ1 | H-Donor | B:SER5:OG | H-Acceptor |
| A:ASN108:HD21 - B:ALA9:O | 2.17069 | Hydrogen Bond | Conventional Hydrogen Bond | A:ASN108:HD21 | H-Donor | B:ALA9:O | H-Acceptor |
| A:ASN108:HD22 - B:LEU8:O | 3.06757 | Hydrogen Bond | Conventional Hydrogen Bond | A:ASN108:HD22 | H-Donor | B:LEU8:O | H-Acceptor |
| A:LYS132:HN - B:SER1:OG | 2.85047 | Hydrogen Bond | Conventional Hydrogen Bond | A:LYS132:HN | H-Donor | B:SER1:OG | H-Acceptor |
| B:SER5:CB - A:GLU75:OE2 | 3.46794 | Hydrogen Bond | Carbon Hydrogen Bond | B:SER5:CB | H-Donor | A:GLU75:OE2 | H-Acceptor |
| B:ALA9:CB - A:PHE74 | 3.86511 | Hydrophobic | Pi-Sigma | B:ALA9:CB | C-H | A:PHE74 | Pi-Orbitals |
| A:LYS71 - B:LEU6 | 4.69039 | Hydrophobic | Alkyl | A:LYS71 | Alkyl | B:LEU6 | Alkyl |
| A:LEU111 - B:LEU8 | 5.20806 | Hydrophobic | Alkyl | A:LEU111 | Alkyl | B:LEU8 | Alkyl |
| B:PRO2 - A:ILE142 | 4.71973 | Hydrophobic | Alkyl | B:PRO2 | Alkyl | A:ILE142 | Alkyl |
| B:ALA9 - A:LEU111 | 4.44657 | Hydrophobic | Alkyl | B:ALA9 | Alkyl | A:LEU111 | Alkyl |

**Table S3. Oligonucleotide primers used in this study.**

| | |
|---|---|
| I. Primers used to generate NLS mutants in GL2 and PDF2. Codons representing the Ala substitutions are given in bold. | |
| gl2_L22_24AA_F | **GCA** TCT **GCC** GCT GGG ATA TTC |
| gl2_L26A_R | AGA **GGC** GGC TGG AGA GG |
| gl2_L705A_L707A_F | **GCC** GCC **GCC** CAA ACC CTC ATC AAC |
| gl2_L702A_L703A_R | TGT **TGC GGC** CGA CCC TCC TTG GCT G |
| gl2_EAR_N_Del_F | GCT GGG ATA TTC CGG AAT GC |
| gl2_EAR_N_Del_R | GGC TGG AGA GGA GAA AAA GTC |
| II. Primers used for DNA assembly to clone GL2 and mutants in SR54 binary vector. | |
| SR54_EYFP:GL2_F | TCAAGCTTCGAATTCTGCAGTCGAC ATG TCA ATG GCC GTC GAC ATG |
| SR54_p1300_GL2_R | CCACCGCGGTGGAGCTCG TCA TTA GCA ATC TTC GAT TTG |
| III. Primers used for pENTR-dTOPO directional cloning. The coding sequences are shown in blue and stop codons are indicated in red. | |
| TPL_F | CACC ATG TCT TCT CTT AGT AG |
| TPL_R | TCA TCT CTG AGG CTG ATC AG |
| Topo_TPR1_F | CACC ATG TCT TCT CTG AGC AG |
| Topo_TPR1_R | TCA TCT CTG AGG CTG GTC AG |
| IV. Primers for genotyping TPL/TPR mutants. | |
| LBb1.3 | ATTTTGCCGATTTCGGAAC |
| TPL_LP | AGACCCTGCTTCTAGGTGTG |
| TPL_RP | TTTGTTCATGTACCTGGAGGC |
| TPR2-2 LP | TCAGCATCAAAGACTGCAATG |
| TPR2-2 RP | TGGGAAGGTGATTCGTTGTAC |
| TPR3-1LP | GTTCTCTTGCAGCCTCAATTG |
| TPR3-1RP | TTCCCACAATGTGATTTCTCC |
| TPR4-1 GTF | ATGTCGTCACTCAGCAGAGAACTC |
| TPR4-1 GTR | GCAAAGCTGATGTTGCCAGTTCAA |
| TPR1-Geno_F | AACCAGACCTATGGGAATC |
| TPR1-Geno_R1 | CCACCGTGATAAGAGTATAGC |
| TPR1-Geno_R2 | CATCCCAGACCTTAATTCAC |
| V. Oligonucleotide used for construction of SR54 EYFP:SRDX:gl2$^{EAR_N}$ by DNA assembly. SRDX encoded residues are indicated in bold. The EYFP and GL2 sequences are shown as magenta and blue, respectively. | |
| SRDX | CT CAA GCT TCG AAT TCT GCA **CTG GAT CTG GAT CTG GAA CTG CGC CTG GGC TTT GCG** ATG TCA ATG GCC GTC GAC ATG |

### SI References

**Brady, S.M., Orlando, D.A., Lee, J.Y., Wang, J.Y., Koch, J., Dinneny, J.R., Mace, D., Ohler, U., and Benfey, P.N.** (2007). A high-resolution root spatiotemporal map reveals dominant expression patterns. Science **318,** 801-806.

**Gilding, E.K., and Marks, M.D.** (2010). Analysis of purified glabra3-shapeshifter trichomes reveals a role for NOECK in regulating early trichome morphogenic events. Plant J **64,** 304-317.

**Le, B.H., Cheng, C., Bui, A.Q., Wagmaister, J.A., Henry, K.F., Pelletier, J., Kwong, L., Belmonte, M., Kirkbride, R., Horvath, S., Drews, G.N., Fischer, R.L., Okamuro, J.K., Harada, J.J., and Goldberg, R.B.** (2010). Global analysis of gene activity during Arabidopsis seed development and identification of seed-specific transcription factors. Proc Natl Acad Sci U S A **107,** 8063-8070.

**Marks, M.D., Wenger, J.P., Gilding, E., Jilk, R., and Dixon, R.A.** (2009). Transcriptome analysis of Arabidopsis wild-type and gl3-sst sim trichomes identifies four additional genes required for trichome development. Mol Plant **2,** 803-822.
